# A third type of PETase from the marine *Halopseudomonas* lineage

**DOI:** 10.1101/2024.12.31.630877

**Authors:** Onur Turak, Andreas Gagsteiger, Ashank Upadhyay, Mark Kriegel, Peter Salein, Seema Agarwal, Erik Borchert, Birte Höcker

## Abstract

The enzymatic degradation of polyethylene terephthalate (PET) offers a sustainable solution for PET recycling. Over the past two decades, more than 100 PETases have been characterized, primarily exhibiting similar sequences and structures. Here, we report new PET-degrading α/β hydrolases, including *Halo*PETase1 from the marine *Halopseudomonas* lineage, thereby extending the narrow sequence space by novel features at the active site. The crystal structure of *Halo*PETase1 was determined to a resolution of 1.16 Å, revealing a unique active site architecture and a lack of the canonical π-stacking clamp found in PETases so far. Further, variations in active site composition and loop structures were observed. Additionally, we found five more enzymes from the same lineage, two of which have a high similarity to type IIa bacterial PETases, while the other three resemble *Halo*PETase1. All these enzymes exhibited high salt tolerance ranging from 2.5 to 5 M NaCl leading to higher total product releases upon PET degradation at 40 or 50 °C. Based on these findings, we propose an extension of the existing PETase classification system to include type III PETases.

## Introduction

The recycling of commodity plastics is currently under extensive investigation, mainly motivated by the need to mitigate environmental plastic pollution and to reduce unsustainable resource depletion^1^. This is supported by recent efforts toward a legally binding UN plastic treaty, which emphasizes plastic recycling as a key component of a holistic approach to mitigating plastic pollution^2^. However, only 9 % of plastic waste is globally recycled^3^. This indicates the need for improvement of the current recycling infrastructure for the implementation of a circular economy for plastics.

Polyethylene terephthalate (PET) is an aromatic polyester and one of the commodity plastics that is widely recycled in comparison to other plastics, such as polypropylene, polyurethane or polyvinyl chloride^4^. The use of enzymes for the recycling of PET can provide an almost complete conversion to monomeric starting materials such as terephthalic acid (TPA)^5–7^. This product can be incorporated into the production of new PET (recycling)^8^ or the synthesis of new and valuable chemicals such as vanilline (upcycling)^9^. Enzymatic recycling enables the selective de-polymerization of PET without losing material properties (downcycling), which can be the case for mechanical recycling, and requires generally less harsh reaction conditions, e.g. milder temperatures and aqueous buffers, in contrast to chemical recycling^10^.

Enzymes such as improved variants of leaf-branch cutinase (LCC)^6^ and polyester hydrolase 7 (PHL7^11,12^, also called PES-H1^7^) are potential candidates for their application in industrial PET recycling^13,14^. These enzymes de-polymerize low crystalline PET at the glass transition temperature (65-75 °C) quite efficiently^15^. However, most of the post-consumer PET (pcPET) waste has a crystallinity up to 45 % leading to drastic reduction of the enzymatic rates^15,16^. Contaminants, additives and polymer blends as well as the size of the plastic can affect the same, and thus, the energy and water-demanding cleaning, sorting and pre-treatment of pcPET for crystallinity reduction is key for efficient enzymatic de-polymerization^17,18^. Current PETases are already quite optimized and show relatively high de-polymerization rates at the glass transition temperature, but the challenges remain. This leaves the future role of enzymes for PET recycling undefined. Novel enzymes with for example, the ability to degrade high crystalline PET or a tolerance against a wide range of reaction conditions, could enhance the sustainability of PET recycling^19^ and have an impact on the future role of enzymes in PET recycling as a part of a circular plastic economy.

Here in this work, we apply a sequence homology search in marine metagenomes from the Tara Oceans project^20^ with the goal of identifying more versatile enzymes. Several studies indicated that marine-derived enzymes can be highly salt-, pressure-, cold-, and organic solvent tolerant^21–24^. Most recently, PET-degrading enzymes with a NaCl tolerance up to 5 M were identified from the deep-sea, including areas such as hydrothermal vents^25^. Due to a stability at lower water activity in the presence of high ionic strength or at low temperatures, these enzymes are more likely to show organic solvent tolerance^26^. The “crystallinity problem” of pcPET could be circumvented in a scenario, where the PET is dissolved in benign co-solvent systems^27^. Taken together, marine-derived PETases can help to build additional and alternative routes for the enzymatic recycling of PET.

The need for more versatile enzymes mirrors the need for an expansion of PETase sequence and structure spaces^28^. Over the past 10 years, more than 100 PETases, i.e. PET-degrading α/β hydrolases, were identified and biochemically characterized (see also PAZy database^29^). These enzymes originate mostly from bacteria such as *Pseudomonadota*, *Actinomycetota*, and *Bacillota.* Since these enzymes were mainly identified by sequence homology searches partially based on already identified PETases^30^, they exhibit similar sequence and structure properties. Later enzymes with more diversity were found. These PETases showed to some extent higher thermostabilities^31^ or originated from archaea instead of bacteria^32^.

Although bacterial PETases show higher similarities, they can be distinguished based on some structure and sequence differences^33^. The following criteria are used for categorization: the occurrence and position of disulfide bonds, the amino acid-composition at the active sites (subsite I and II), and the occurrence of an extended loop flanking the active site. A representative of type I enzymes is LCC. These PETases show one disulfide bond at a conserved position and have no extended loop region surrounding their active site. Type II enzymes have two conserved disulfide bonds and show an extended loop region. Representatives are commonly mesophilic enzymes in contrast to PETases from type I. Type II is further subdivided into type IIa and IIb, whereby these two differ in their amino-acid composition in subsite II and the extended loop region. The benchmark *Is*PETase is classified as a type IIb PETase and consists of a stabilizing disulfide bond close to its active site. A representative of type IIa PETases is polyester hydrolase (PE-H) from the marine bacterium *Pseudomonas aestusnigri*^34^. This bacterium belongs to the bacterial lineage *Halopseudomonas* (formerly *Pseudomonas pertucinogena*^35^), which has been a good source for the identification of PET-degrading enzymes^36^, such as PE-H from *H. aestusnigri* (*Haes*_PE-H)^34^, PE-H from *H. formosensis* (*Hfor*_PE-H)^37^, or *Pbauz*Cut from *H. bauzanensis*^38^. Bioinformatic analysis further suggested putative PETases in *P. sabulinigri*, *P. pachastrellae* and *P. litoralis*^33^. All of these are classified as type IIa PETases.

In this work, we identify and characterize six novel PETases from the bacterial genus *Halopseudomonas*. We show that four enzymes (*Halo*PETase1, 2, 3, and 4) cannot be categorized into one of the known PETase types. Therefore, we establish together with the known cutinase from *Pseudomonas mendocina* (*Pm*C) the new bacterial PETase type III and explore by bioinformatic analyses more homologs of this type in the proteome of *Halopseudomonas*. In addition, we test the so far unexplored salt-tolerance of PETases originating from *Halopseudomonas*. These enzymes show improved PET-degradation in the presence of 2.5 to 5 M NaCl. Thus, we (1) provide novel enzymes for the PET-degradation in a high-salt environment. This might enable the use of these enzymes under more chemically demanding conditions for PET waste management. (2) We explore and highlight the potential of the *Halopseudomonas* lineage as a source for PETases or other biotechnologically useful polyesterases. And (3), we extend the narrow sequence space of PETases and enable the search for novel enzymes by covering more diverse candidates.

## Results and Discussion Screening for novel PETases

To identify novel PETases from the marine environment, a profile hidden Markov-Model (pHMM) search was applied towards marine bacterial metagenomes, published by the Tara Oceans project^20^. 20 target proteins that showed a bit-score higher than 100 were selected to analyze their ability for PET degradation. These proteins were recombinantly produced in *E. coli*. Subsequently, purified proteins were spotted onto PBS-agar plates supplemented with either BHET, PET-NP or PCL to characterize their enzymatic activity towards these substrates.

We observed the formation of zones of clearances (halos) for 14 target proteins towards all substrates (Figure S1 and Table S1). Three hits, PSW62-1, *Halo*PETase1 (screening name: PSW62- 2) and ASW29-1 indicated halos on PET-plates after seven days of incubation at 30 °C. We analyzed their affiliation and identified the origin of their gene sequences in metagenomes associated with *Pseudomonas sp.* PSW62-1 and ASW29-1 have a sequence identity of 89.2 % with the PET-active polyester hydrolase (PE-H) from *Halopseudomonas aestusnigri*^36^. In the case of *Halo*PETase1, a cutinase from *Pseudomonas mendocina* (*Pm*C) is the closest related and characterized PETase^39^. Due to a sequence identity of 62.4 % with *Pm*C and sequence identities of less than 34 % to the next related and characterized PETases (see Table S2), we concluded that *Halo*PETase1 is a promising candidate for extension of the narrow sequence space of currently known PETases.

## Quantification and optimization of PET-degradation by *Halo*PETase1 (PSW62-2)

*Halo*PETase1 was produced and tested in the functional agar screening with an N-10xHis-SUMO-tag. For subsequent experiments the tag was removed from the enzyme during purification (see also Figure S2 for SDS-gel). We investigated the PET degradation (Figure 1) conditions by *Halo*PETase1 first using PET-coated well-plates^40^ and then using in solution immersed PET-films to assess the potential of this enzyme.

**Figure 1:**
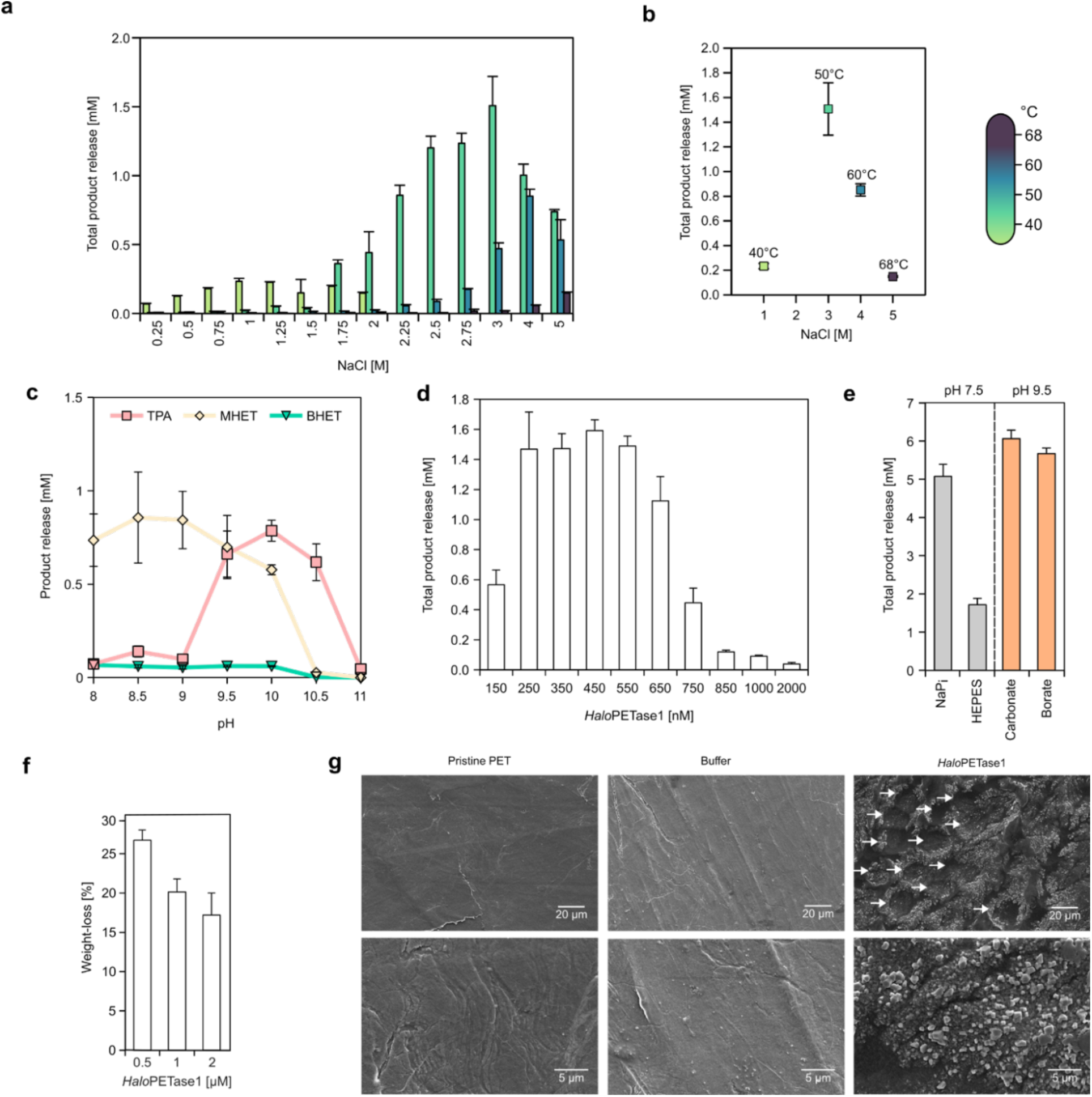
Reaction conditions for PET-degradation by *Halo*PETase1 (PSW62-2). **a)** NaCl-dependent total product release (sum of BHET, MHET, and TPA) at 40, 50, 60, and 68 °C in 20 mM NaPi, pH 7.4. **b)** NaCl optimum concentrations for PET degradation shift upon increase of reaction temperature. **c)** pH-dependent PET-degradation, using 20 mM Bicine for pH 8 and 8.5, 20 mM CAPSO for pH 9 or 9.5, 20 mM CAPS for pH 10, 10.5 or 11. Incubation at 50 °C for 20 h. **d)** Enzyme concentration dependent PET degradation at 50 °C, 3 M NaCl and pH 7.4 for 20 h. **e)** pH- and buffer-dependent PET-degradation. 0.1 M buffer, 3 M NaCl, 0.5 µM *Halo*PETase1. Analyses in **a-e** were performed with the PET-coated well-plate assay^40^. **f)** Weight-loss analysis of solution immersed PET-films upon buffer and *Halo*PETase1 solution treatment for 7 days. **g)** SEM micrographs of untreated (i.e. pristine), buffer and *Halo*PETase1 solution treated films from experiment shown in **f**. Pit-hole like formations upon degradation by *Halo*PETase1 are indicated by arrows. Enzymatic degradation was performed with 3 technical replicas, error bars indicate standard deviations.

*Halo*PETase1 was found in marine metagenomes and includes an N-terminal signal peptide indicating that the protein is likely secreted into the saline marine environment. Therefore, we tested degradation of PET using a NaCl gradient ranging from 0.25 to 2 M at 40, 50, and 60 °C (Figure 1a). The highest total product release at 40 °C after 20 h was observed in the enzyme solution with 1 M NaCl (Figure 1b), whereby higher NaCl concentrations led to a decreased product release. Since the total product release increased continuously from 0.25 to 2 M NaCl at 50 °C, we assumed that NaCl concentrations higher than 2 M will lead to even higher total product releases. Consequently, we also tested PET-degradation at 50, 60 and 68 °C with up to 5 M NaCl. Notably, the optimal NaCl concentrations increased with higher reaction temperatures (Figure 1b). We observed the highest total product release at 50 °C with 3 M NaCl. At 60 °C, it decreased in comparison to the PET degradation at 50 °C, but the NaCl optimum increased to 4 M. Similarly, at 60 °C the total product release was even lower than at 50 °C, but the NaCl optimum increased even up to 5 M.

To test the effect of NaCl on the enzymatic thermostability, we incubated *Halo*PETase1 up to three days at 50, 60, and 68 °C with either 1, 3, or 5 M NaCl. Subsequently, the residual activity was measured by a *para*-Nitrophenol butyrate (*p*NPB) assay (Figure S3). The residual activity after 72 h at 50 °C with 5 M NaCl apparently did not decrease. However, we observed a reduction of the residual activity to 49 % and 74 % with 1 and 3 M NaCl after 72 h, respectively. At 60 °C, the residual activity dropped to 0.6 % with 1 M NaCl after 4 days, while with NaCl concentrations of 3 and 5 M it decreased to 2.7 % and 15.3 %, respectively, after 72 h at the same temperature. Finally at 68 °C, no more residual activity could be observed for all NaCl concentrations after 1 h. Taken together, these results indicate a stabilizing effect of high NaCl concentrations on *Halo*PETase1 at higher temperatures.

Next, we investigated the pH dependent degradation in the PET-coated well-plate setup with 100 nM *Halo*PETase1. We observed that the highest total product release at 50 °C with 3 M NaCl occurs at pH values between 9.5 and 10, as indicated by the sum of TPA, MHET and BHET concentrations (Figure 1c). The degradation of PET with sodium phosphate buffer at pH 8 mainly led to the formation of MHET. However, the degradation of PET with CAPS buffer at pH 10.5 mainly led to the formation of TPA. Both conditions, nevertheless, indicated similar total product releases.

After identifying the apparent optimal enzyme concentration within this experimental framework (Figure 1d), we performed buffer and pH-dependent PET-degradation experiments with 500 nM *Halo*PETase1. The release of acidic products, such as MHET or TPA, during enzymatic hydrolysis of PET can reduce the pH if not sufficiently buffered. This can affect the enzymatic rates due to effects on protein stability and activity. Further, the choice of buffer can have a concentration dependent effect on enzyme rates of PETases as well^41^. Thus, we increased the buffer concentrations to 0.1 M to gain a higher total product release. When degradation was tested at pH 7.5 with sodium phosphate or pH 9.5 with carbonate or borate buffer, total product releases increased yielding between 5 to 6 mM (Figure 1e). In the case of 0.1 M HEPES buffer at pH 7.5, less than 2 mM total product release was observed, indicating an inhibiting effect of this buffer.

Further, we analyzed PET-degradation by *Halo*PETase1 using medium crystallinity PET-films (∼15 % crystallinity [see also Supplementary Figure S10], 150-200 µm thickness, 1 cm x 1 cm, ∼40 mg) in a weight-loss experiment. Here, PET films were immersed in an enzyme solution for seven days at 50 °C, and weight loss of the films was subsequently measured (Figure 1f). The highest weight-loss (∼25 %) was observed with 0.5 µM *Halo*PETase1. An increase of the enzyme concentration to 1 and 2 µM showed lower weight-losses of about 20 and 17 %, respectively. Correspondingly, scanning electron microscopy (SEM) micrographs were acquired to analyze changes in the surface topography of the PET films after immersion in the enzyme solution (Figure 1g). In comparison to the pristine and negative control (only buffer treated) PET films, the surface of 0.5 µM *Halo*PETase1 treated film showed the formation of pit-hole like topographies with diameters ranging from 10 to 20 µm. Similar topography has been observed previously for the thermostable PETase PHL7, where larger pits were seen on PET-disks with a similar crystallinity (15.8 %) upon treatment^42^. Notably, crystal-like formation on the treated film surfaces may indicate NaCl crystals due to buffer conditions.

Taken together, these results demonstrate that *Halo*PETase1 degrades PET in both the PET-coated well-plate assay and the solution immersed PET-film assay. *Halo*PETase1 is a salt-tolerant enzyme and depends on higher salt-concentrations for the degradation of PET at elevated temperatures.

## Structure and sequence classification of *Halo*PETase1

Bacterial PETases can be classified as type I, type IIa, and type IIb based on their structural and sequence features near their active sites^12,33^, such as the presence of an extended loop surrounding the active site, the number and position of disulfide bonds, and the amino acid composition at subsites I and II. Type I PETases have no extended loop regions and show one disulfide bond. In contrast, type II PETases have extended loop regions and two disulfide bonds; they are further subdivided into type IIa and IIb based on differences in the extended loop regions.

To better classify *Halo*PETase1, we elucidated its X-ray crystal structure to a resolution of 1.16 Å (PDB code: 9hl5, Table S3) and compared it to existing PETases (Figure 2). *Halo*PETase1 has the canonical α/β hydrolase fold illustrating highly conserved spatial positions for its catalytic residues S156, D206, and H236 as known for *Is*PETase (type IIb). There are two disulfide bonds observed: Db I (C64-C126) and Db II (C273-C276)(Figure 2a). While two bonds are characteristic for type II PETases, these are, however, located at different positions in *Halo*PETase1. Notably, we could not observe a disulfide bond flanking the active site, as it is known for *Is*PETase^33^.

**Figure 2:**
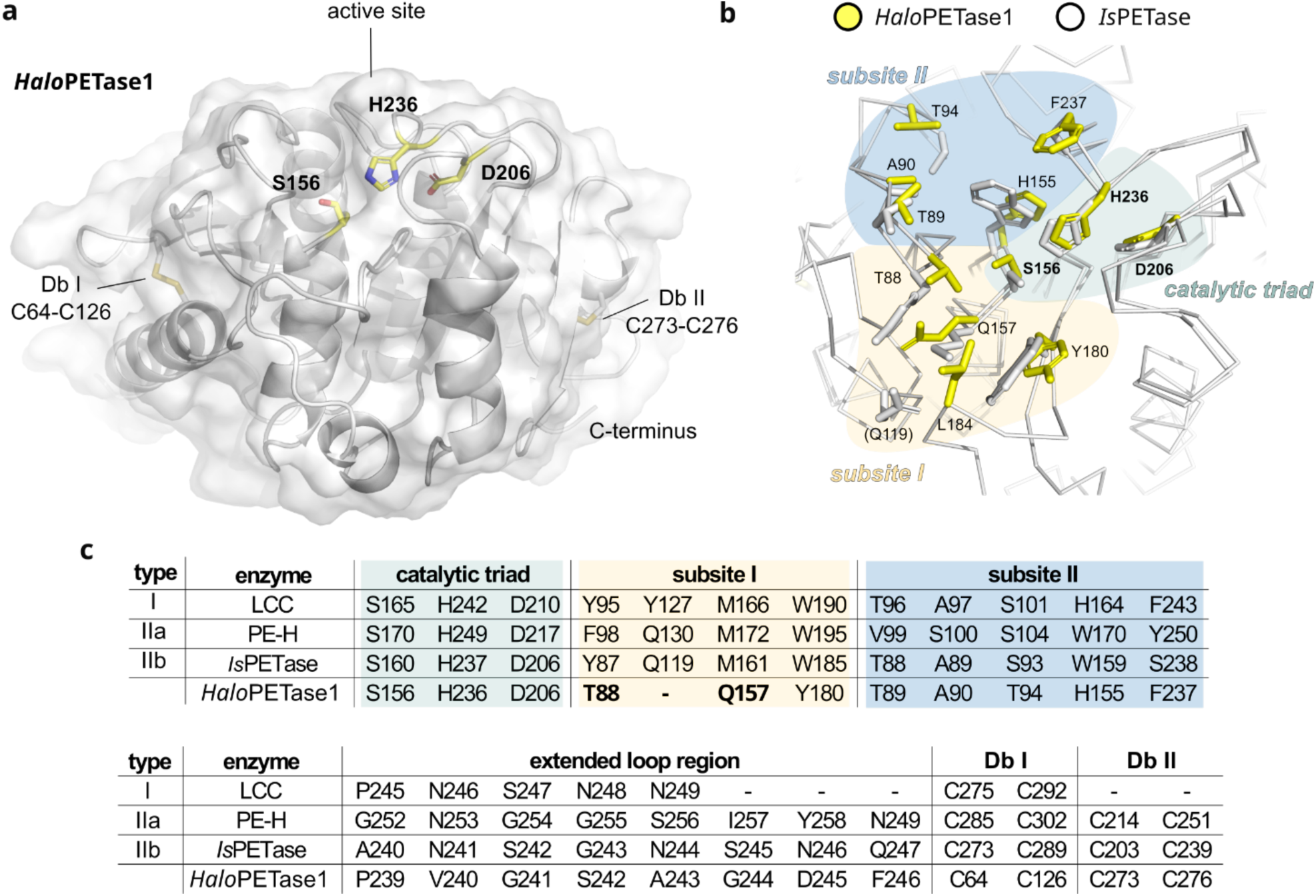
Structure and sequence features of *Halo*PETase1 in comparison to previously characterized PETases. **a)** Crystal structure of *Halo*PETase1 with a resolution of 1.16 Å (PDB code: 9hl5). Cartoon representation of the protein backbone overlayed with a surface representation. The catalytic residues at the active site S156, H236 and D206 are labeled in bold. These residues and the disulfide bonds, C64-C126 and C273-C276, are shown as sticks. **b)** Superimposed active sites of *Is*PETase (PDB: 5xjh) and *Halo*PETase1. Ribbon representation of the backbones and stick for sidechains that are associated with PET-activity. Respective amino acids for *Halo*PETase1 are shown in yellow with corresponding labels. These were inferred from the alignment with *Is*PETase and according to Joo *et al.*^33^. Residues from *Is*PETase are shown as grey sticks. Only residue Q119 from subsite I of *Is*PETase is shown in brackets as no equivalent residue could be derived for *Halo*PETase1. **c)** Amino acid occupation at the catalytic site, subsites I and II, the extended loop region, and the presence of one or two disulfide bonds of representative PETases from type I (LCC), IIa (PE- H), IIb (*Is*PETase) and *Halo*PETase1. The catalytic triad, subsites I and II are highlighted by the same colors as in **b**, thus corresponding to these structure sites.

Previous structure analyses on the binding mode of PET substrate modules to PETases suggested an aromatic interaction between the benzene moiety of the substrate with two opposing aromatic amino acids at the active site. For example, PHL7 (type I) indicated a distorted T-shaped π-π-interaction of F63 and W156 with the benzene moiety of TPA in a co-crystal structure^12^. Similarly, *Is*PETase showed, depending on the study, up to two π-π interactions by Y87 and W185 (docking of tetrameric PET module and mutational analysis)^33^ or W185 (QM/MM analysis or complex crystal structure with HEMT)^43,44^. Altogether, type I and II PETases consist of two opposing aromatic amino acids such as Y/F and W at equivalent positions to F63 and W156 in PHL7, and Y87 and W185 in *Is*PETase.

In *Halo*PETase1 on the other hand we observed the non-conserved aromatic acid Y180 in subsite I (equivalent to W185 in *Is*PETase, Figure 2b and c). However, instead of an aromatic residue at the equivalent position of Y87 in *Is*PETase, we found T88 for *Halo*PETase1. Thus, the so-called π- stacking “clamp” does not exist for *Halo*PETase1. Further, we hypothesize that the backbone amide protons of T88 and Q157 are responsible for the formation of the oxyanion hole in *Halo*PETase1 since these residues are located at equivalent positions as M161 and Y87 in *Is*PETase. With the goal of restoring the otherwise conserved π-stacking clamp we tested single mutations by substituting T88 with either T88W, T88F, or T88Y. These mutations led to a reduction of total product release by almost half of the wild-type enzyme (Figure S4a). A similar result was observed for the Q157M mutation. Here, we intended to restore the highly conserved methionine and reduce the polarity in the active site. Neither the restoration of the π-stacking clamp nor the reintroduction of the highly conserved methionine showed improved total product releases. We assume that other factors may play a role. For example, larger structural changes affecting enzymatic activity or reduction in thermostability might be responsible for the lower product releases under these reaction conditions. The restoration of the π-stacking clamp might also require further mutations for a co-operative effect to enable the correct architecture for a π-stacking clamp. Amino acids in the vicinity of the introduced aromatic residues might not pair well due to unfavorable interactions or steric hindrances. For example, residue L163 of *Halo*PETase1 is apparently placed between the putative π-stacking clamp, possibly blocking the insertion of the substrate between the aromatic residues (Figure S4b). It should be noted that this residue is placed on a unique and extended loop which is not observed for *Is*PETase. Due to the existence of this loop, no equivalent residue for Q157 in *Is*PETase could be derived for *Halo*PETase1.

The extended loop region, which is known for type II PETases, is present in *Halo*PETase1 as well. However, only P239 and G241 in *Halo*PETase1 are also present at equivalent positions in LCC (type I) and PE-H (type IIa).

We found similarities to other PETase types in subsite II of *Halo*PETase1. The amino acid occupation at this part of the active site resembles the one of LCC (type I). The only difference is the presence of T94, which is S101 in the case of LCC (type I). A serine seems to be conserved at this position in other PETases: S104 in PE-H (type IIa) or S93 in *Is*PETase (type IIb).

## A characteristic active site architecture of *Halo*PETase1 and homologs

The structural alignment of *Halo*PETase1 and *Is*PETase at subsite I indicate that Q119 (*Is*PETase) is located on a loop which is distinct from the corresponding loop of *Halo*PETase1 (Figure 2b and c). Therefore, no corresponding residue could be derived for *Halo*PETase1. This difference was also observed for spatially similar loops from representative PETases of other types, such as LCC (type I, Y129) and PE-H (type IIa, Q130) (Figure 3a). In contrast, *Halo*PETase1 has a distinct and extended loop (loop 3) due to an insertion of five amino acids (Figure 4a and b). Notably, L163 in *Halo*PETase1, not observed for other PETases, is located on this loop and is potentially involved in substrate binding.

**Figure 3:**
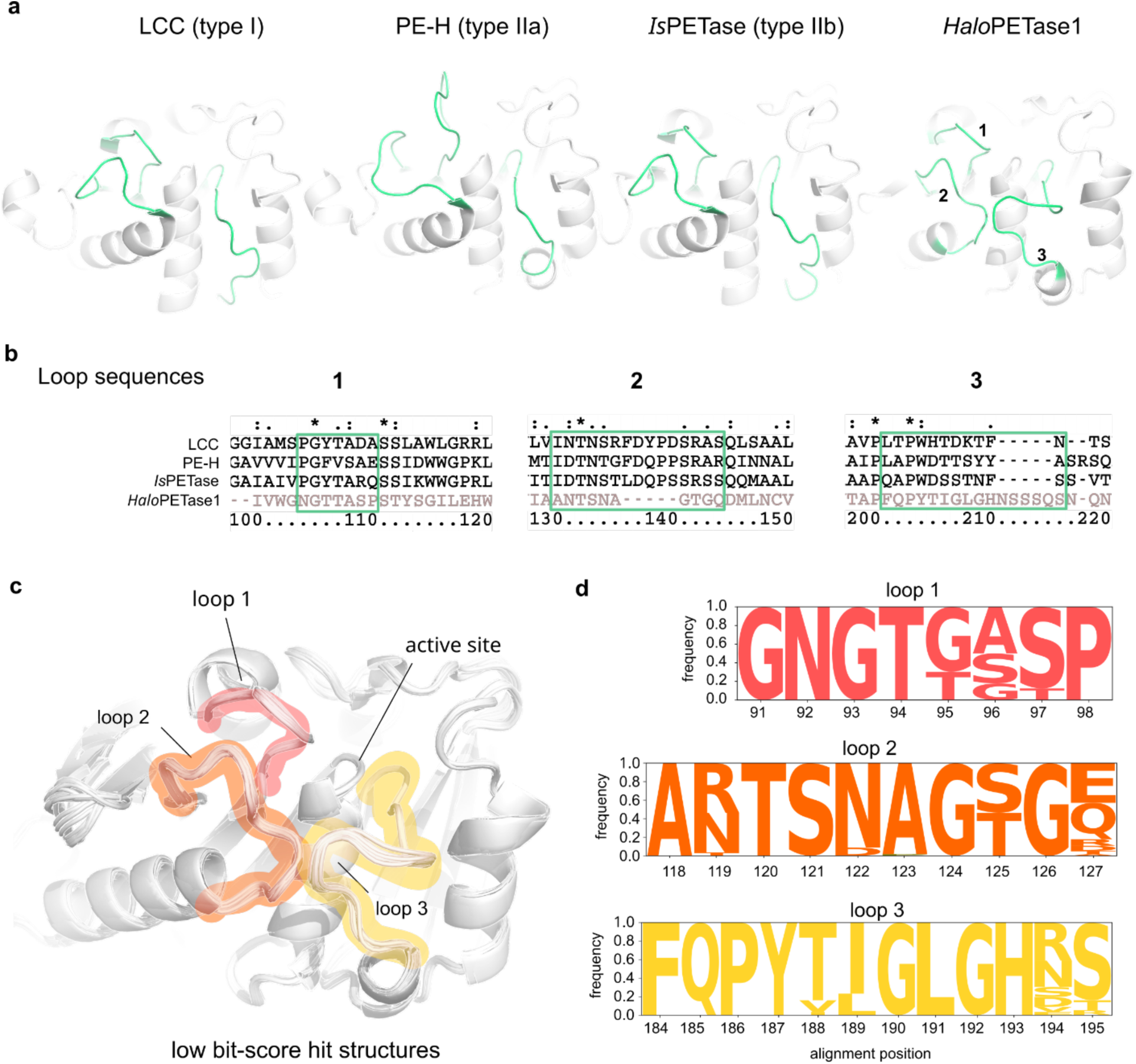
Loops surrounding the active sites in representative PETases from type I, IIa, IIb, and *Halo*PETases. **a)** Loops of interest in crystal structures of LCC (PDB: 4eb0), *Haes*_PE-H (PDB: 6sbn), *Is*PETase (PDB: 5xjh) and *Halo*PETase1 (PDB: 9hl5) are colored green. These loops are numbered for *Halo*PETase1. **b)** Corresponding sequence alignments for loop regions in **a**. **c)** Aligned ColabFold^45^ model structures of low bit-score hits from *Halopseudomonas* proteome including *Halo*PETase2, 3, and 4. Structure alignment was performed globally with *align* in PyMOL over backbone C_α_ atoms (5 refinement cycles, >2 Å outlier rejection) without the first 40 residues to exclude the N-terminal signal peptides. Corresponding loops are highlighted in red (loop 1), orange (loop 2), and yellow (loop 3). **d)** Logo plot of aligned loop regions for low bit-score hits.

**Figure 4:**
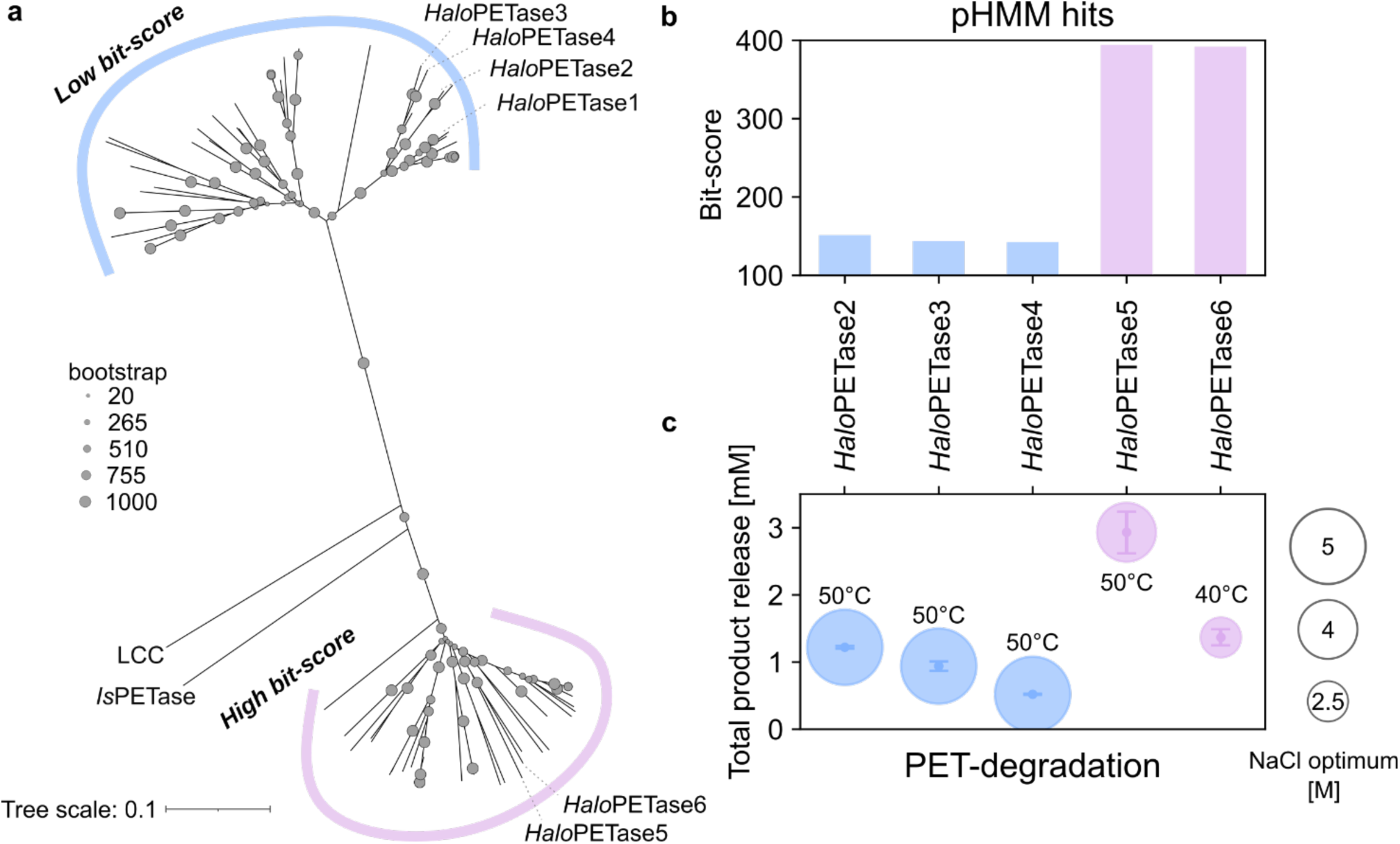
Phylogenetic tree of putative PETases from *Halo*PETase1 homologs in the proteome of *Halopseudomonas*. **a)** Neighborhood-joining tree of all pHMM hits in the proteome of *Halopseudomonas* separated into low bit-score group and high bit-score group sub-clades. **b)** Bit-scores of selected *Halo*PETases2 to 6. **c)** Total product releases of respective *Halo*PETases at their apparent NaCl- and temperature-optimum. The PET-degradation was performed with PET-coated well plates for 20 h and soluble products were analyzed via UHPLC. Enzymatic degradation studies were performed with 3 technical replica, error bars indicate standard deviations.

Another characteristic loop (loop 2) was observed, close to loop 3, where in other PETases a residue of subsite I is present, e.g. Q119 of *Is*PETase or Y95 of LCC. Overall, loop 2 is shorter than corresponding loops of other PETases by five amino acids (Figure 4b). There is potentially no residue on loop 2 of *Halo*PETase1 directly involved in substrate binding. Nevertheless, loop 2 flanks the active site and is in close contact with loop 1 and 3, indicating an indirect effect on substrate binding. Finally, loop 1 of *Halo*PETase1 shows a similar amino acid composition and has the same length as *Is*PETase and LCC. However, a major difference is the missing aromatic residue for the π stacking interaction with the substrate. Note that loop 1 in PE-H is longer than in other PETases^34^.

Based on these observations we performed another pHMM search in the proteome of *Halopseudomonas* with a profile consisting of already biochemically characterized PETases. The goal was to identify more PETases resembling *Halo*PETase1. We identified hit sequences with bit- scores ranging from 125.2 to 412.4 (Table S4). The hits were separated into a low bit score [125.2-200) and a high bit-score [200-412.4] group.

Upon modeling the 3D-structures of *Halo*PETase1 homologs from the low bit-score group in ColabFold^45^ we found the same active site backbone architecture (see Figure 3c). Thus, loops directly involved or flanking the active site (loop 1-3) appear conserved. This is also seen in the amino acid composition and frequency of their occurrence in respective loops, as shown in Figure 3d.

In addition to these conserved loops, we found highly conserved amino acids in parts of the active site (logo plot in Figure S9). All amino acids in subsite I among these homologs (orange letters in Figure S9) are strictly conserved (alignment positions T94, Q163, and Y186). Subsite II composition indicates more variance at the following alignment positions: G/T95, A/S/G96, T/G100, H161, and F243. And finally, the extended loop region indicates also more variability: P245, V/A/I246, G/S/N247, N/D/S248, A/G249, G250, D/G/E251, and Y/F252. The known PETase with highest sequence identity to *Halo*PETase1 (62.4 % identity, see Table S2), *Pm*C, shows also high sequence similarity to low bit-score *Halo*PETases, including *Halo*PETase2, 3, and 4 (MSA in Figure S7).

In conclusion, we observed that conserved features of the existing PETase types I, IIa and IIb were missing in *Halo*PETase1. The subsite I does not consist of a π-stacking clamp and has different amino acids forming the oxyanion hole. Additionally, one amino acid could not be derived from the structural alignment with *Is*PETase, due to the existence of another extended loop at this position. We identified two disulfide bonds as in type II PETases, but these were located at different positions.

## Sequence characterization of *Halo*PETase1 homologs

Construction of an unrooted phylogenetic tree with hit protein sequences (see Table S4) illustrated the separation into two sub-clades, indicating two sequentially distinct groups (see Figure 4a). The hit sequences occurred in one of the two sub-clades according to their assigned bit-scores.

Currently known PETases consist of a lipase-box motif which includes the catalytic serine and has the G-X-S-X-G sequence. For example, type I PETases such as LCC and PHL7 have a G-H-S-M-G^6,12^ and type II PETases such as *Haes*_PE-H (type IIa) and *Is*PETase (type IIb) have a G-W-S-M-G motif^34,46^. The high bit-score group represents the lipase-box motif G-W-S-M-G as other type II PETases. Notably, LCC and *Is*PETase, as indicated in the tree, are closer related to the high bit- score than the low bit-score group. Pairwise sequence alignment identities of *Is*PETase and LCC with all hits from the high and low bit-score groups support this observation: the sequence identity of *Is*PETase and LCC range from 46 to 54 % for high bit-score sequence alignments (Figure S5), whereas the sequence identities for both PETases ranged from 23 to 31 % for low bit-score sequence alignments (Figure S6). Finally, hits in the low bit-score group represent the lipase-box motif G-H-S-Q-G (see Figure S9), which we observed for the PET-degrading enzyme *Pm*C and the newly characterized *Halo*PETase1 (see MSA in Figure S7).

We included *Halo*PETase1 in the phylogenetic tree and observed that it can be found in the sub- clade with other low bit-score hits (Figure 4a). *Halo*PETase1 shows high sequence identities to other hit sequences in the same group, ranging from 63 to 98 % (Figure S5). In contrast, sequence identities in the high bit-score group are much lower, ranging from 26 to 32% (Figure S6).

## *Halo*PETase1 homologs degrade PET

We selected three hit sequences from the low (*Halo*PETase2, 3, 4) and two from the high bit-score clade (*Halo*PETase5, 6) as representative candidates from both bit-score groups for testing their PET hydrolysis activity (see Figure 4b for pHMM bit-scores). The hits from this representative set are all located on one metagenome derived genome of *H. sabulinigri* HyVt376 obtained from a marine hydrothermal vent site sediment^47^ (see boldly highlighted target proteins in Table S4). The selected enzymes were produced by heterologous expression in *E. coli* (Figure S2) and tested for PET-degradation (Figure S8). Like *Halo*PETase1, all enzymes showed hydrolysis activity on PET-coated well plates at high concentrations of NaCl. *Halo*PETase2-6 indicated apparent optima between 2.5 and 5 M NaCl and 40 to 50 °C (see Figure 4c).

## Conclusion

In this study, we present novel and highly halo-tolerant PETases from the bacterial lineage of *Halopseudomonas*. Our bioinformatic analysis shows that there are two PETase groups in this lineage, whereby *Halo*PETases 5 and 6 are more closely related to existing PETases such as LCC and *Is*PETase. More specifically, the sequences of these enzymes resemble type IIa PETases such as *Haes*_PE-H. In contrast, *Halo*PETases1, 2, 3, and 4 form a distinct active site, which is especially seen in the amino acid composition of the subsite I and in the extended loop region. No typical π- stacking clamp could be identified, and an atypical lipase-box motif for bacterial PETases (G-H-S-Q-G) was observed. In addition, there are two disulfide bonds that are located at different positions in comparison to the ones found in other PETase types. Loops surrounding the active site showed either an extended or shortened length compared to known PETases. Therefore, we could not categorize these enzymes according to the widely accepted classification system for PETases^12,33^. We propose an extension of this PETase classification system by type III PETases that include *Halo*PETase1, 2, 3, and 4, as well as *Pm*C.

We believe that the identification of *Halo*PETases enables the discovery of further polyesterases in the bacterial lineage of *Halopseudomonas*, especially PETases of type III. This notion is supported by the discovery of several homologs in the low bit-score group (type III), including the here- characterized *Halo*PETases1, 2, 3, and 4. In addition, we identified further homologs in the high bits- score group (type IIa), which includes PET-degrading enzymes from other studies^34,38^ and the herein presented *Halo*PETase5, and 6. The identification of PETase type III provides new and distinct sequences. Inclusion of these sequences in future homology searches will enable further discovery of distinct PETases with novel features, ultimately helping to improve sustainable PET recycling by enzymes.

## Material and Methods

### Metagenomic mining for putative PETase candidates in marine bacterial metagenomes

To identify novel PETases with higher sequence and structure diversity to known ones in the PAZy database, a profile hidden-Markov model was prepared. Shortly, the profile was generated with biochemically characterized PETases, as reported previously^29^, and a search was performed against marine metagenomes from the Tara Oceans^20^ and other publicly available metagenomes (accession numbers PRJNA454581 and PRJNA901861) with HMMER3^48^. Hit-sequences with a bit-score higher than 100 were selected for further filtering. The filtering was implemented by excluding duplicates and highly similar sequences as well as sequences that did not include a canonical catalytic triad with Ser-His-Asp. Additionally, hits with more diverse bacterial origins in comparison to existing affiliations were chosen. Finally, twenty target proteins were selected for further analysis.

### Molecular cloning and target protein production in *E. coli* MC1061

The corresponding target genes were truncated if N-terminal signal sequences were present. These sequences were identified by SignalP 6.0^49^. Molecular cloning was performed with the fragment-exchange (FX)-cloning methodology, whereby the truncated target genes were extended with recognition sites for the restriction enzyme *Sap*I using the FX-cloning tool (https://www.fxcloning.org/)^50^. This cloning methodology leads to a single amino acid overhang on each end (N-terminal serine and C-terminal alanine) of the final gene product. Finally, all target genes were codon-optimized for heterologous expression in *E. coli* MC1061. The target genes were cloned into the plasmid vectors p1^50^, p2, p3 or p12^51^, leading to the attachment of an N-10xHis-, N-10xHis- Maltose-binding-protein (MBP)-, N-10xHis-Small Ubiquitin-related MOdifier (SUMO)- or C-6xHis-tag, respectively, to the final protein. The positive control, LCC^ICCG^, was cloned into p12 and the negative control, namely a linker sequence of GSGSGS, was cloned into each vector. Successful cloning was tested by colony PCR and subsequent agarose gel electrophoresis. Finally, genes were verified by sequencing if the corresponding target protein showed hydrolytic activity towards PET. Sequencing was also performed for every negative and the positive control.

For heterologous gene expression in *E. coli* MC1061, chemically competent bacteria were transformed with the plasmid constructs. A pre-culture with 3 ml LB and 100 µg/ml ampicillin was prepared by inoculation and subsequent incubation at 37 °C, overnight and 180 rpm. Next, 20 ml 2YT medium with 100 µg/ml ampicillin in a 100 ml Erlenmeyer flask (main culture) was inoculated with 1 % (v/v) of the pre-culture and incubated at 140 rpm, 37 °C for 2-3 h until an OD600 ∼0.6. Then, the main cultures were transferred to 18 °C and gene expression was induced by adding 100 µl of a 20 % (w/v) L-arabinose stock-solution for a final concentration of 0.1 % (v/v). After 20 h, cultures were placed on ice, decanted into pre-cooled 50 ml tubes and centrifuged for 20 min at 4 °C and 3000 xg. Next, cell pellets were resuspended in 10 ml phosphate buffered saline (PBS) with 10 mM imidazole. Cells were disrupted by ultrasonication, and the lysates centrifuged for 30 min at 4 °C and 3000 xg. The cleared cell lysate was used for further purification.

### Target protein purification by Ni-immobilized metal affinity chromatography (Ni-IMAC)

The soluble fractions of the poly-His-tagged target proteins were subjected to a Ni-NTA column purification using a HisPur™ Ni-NTA Spin Purification Kit (0.2 ml, ThermoFisher) according to the manufacturer’s protocol. Finally, three 200 μl elution fractions were collected and stored at 4 °C for a maximum of two days.

### Functional screening on substrate supplemented agar plates

The production of agar plates supplemented with respective substrates was adapted from Chow *et al.*^52^. PBS agar was prepared with 0.1 M PBS and 1.5 % (w/v) agar, sterilized by autoclaving and allowed to cool down to ca. 60 °C until substrate solutions were added. 20 ml of substrate PBS agar were dispensed into Petri dishes and allowed to solidify under sterile conditions.

### Production of PET nanoparticle (PET-NP) suspension and agar-plates

A stock PET-NP suspension was prepared by dissolving 0.1 g thin pieces of a PET-film (Goodfellow) in 10 ml 1,1,1,3,3,3-hexafluoro-2-propanol (HFIP, Sigma-Aldrich) in a separatory funnel in the fume hood. The solution was dropped slowly into 100 ml pre-cooled and de-ionized water in a glass beaker under vigorous mixing with an Ultra-Turrax™ T25 basic (IKA Labortechnik). Residual HFIP was allowed to evaporate for several days. The suspension (ca. 0.3 mg/ml) was added to 200 ml molten PBS-agar. In all PET-NP suspension preparations, precipitation of PET was observed, which potentially decreased the final concentration of PET-NPs after filtration in the agar plates. Agar-plates with sufficient turbidity after visual inspection were used for further experiments.

### Production of BHET solution and agar-plates

A 1 M stock solution of Bis(hydroxyethyl) terephthalate (BHET, Sigma-Aldrich) was prepared in DMSO. BHET containing PBS-agar plates were prepared with a final concentration of 0.5 M BHET.

### Production of PCL solution and agar-plates

0.25 g of PCL granules (Sigma-Aldrich) were dissolved in 15 ml acetone at 60 °C. This solution was transferred into 500 ml of PBS-agar.

### Functional screening of target proteins

20 μl of Ni-NTA purified target protein were spotted onto PCL, BHET or PET-NP supplemented PBS-agar plates. The plates were incubated at 30 °C for 7 days. Enzymatic hydrolysis activity was detected by the formation of zones of clearances (halos), which became visible due to the decrease of turbidity on the plate.

### Heterologous expression and purification of *Halo*PETase1 (PSW62-2) and single mutants

The codon-optimized gene sequence (without N-terminal signal peptide) of *Halo*PETase1 (screening name: PSW62-2; NCBI accession ID: MAB42154.1) was cloned into the p3 vector, which leads to a® protein with N-10xHis-SUMO-tag upon expression. Chemically competent *E. coli* Shuffle T7 (NEB) were transformed with the respective construct and cultivated on LB-agar plates with 100 µg/ml ampicillin. A pre-culture was prepared in LB with 100 µg/ml ampicillin by cultivation over night at 37°C and 180 rpm. A 2 L culture with TB medium and 100 µg/ml ampicillin in a 5 L Erlenmeyer flask was inoculated with 1 % (v/v) of the pre-culture and grown until an OD600 of ∼0.6 at 37 °C and 140 rpm. The culture was placed at 18 °C and gene expression induced by adding 0.2 % (v/v) from a 20 % (w/v) L-Arabinose stock. Bacteria were harvested after 20 h by centrifugation, the cell pellet was washed in 20 ml PBS and subsequently frozen at -20 °C until further use.

For purification the cell pellets were thawed on ice and resuspended in 40 ml of cold buffer A (50 mM NaPi, 50 mM imidazole, 500 mM NaCl, pH 7.4). Cells were lysed by sonication and the supernatant collected after centrifugation was subjected to a Ni-IMAC with a 5 ml HisTrap^®^ FF column (Cytiva) attached to an ÄKTA pure™ (Cytiva) purification system. Elution was performed by an imidazole gradient with buffer B (50 mM NaPi, 500 mM imidazole, 500 mM NaCl, pH 7.4) ranging from 0 – 50 % over 20 CV.

To cleave the N-10xHis-SUMO-tag from the target protein, 2 mg of SUMO-tag protease SenP2 was added to approx. 40 mg protein (1:20 mass ratio) in the pooled Ni-IMAC fractions. The protein was dialyzed overnight at 4 °C against buffer C (20 mM NaPi, 150 mM NaCl, pH 7.4) to remove imidazole and then further purified by reverse Ni-IMAC. Finally, the target protein was further purified by a size-exclusion chromatography (SEC) using a HiLoad 26/600 Superdex 75 pg column attached to an ÄKTA pure™ (Cytiva) purification system. SEC fractions were pooled, and the protein was concentrated to 10-50 mg/ml. 25-50 µl samples were flash frozen in liquid nitrogen and stored at - 20 °C until further use.

### PET-degradation endpoint measurements in PET-coated well plates and soluble product analysis

96-well plates (Thermo Scientific™ Nunc MicroWell™ flat bottom) were coated with a PET-film according to Weigert *et al*.^40^ 50 µl of an enzyme solution were added per well and the plates were sealed with a gas-tight self-adhesive film. In general, plates with enzyme solutions were incubated at the target temperature for 20 h. Subsequently, all samples were prepared and analyzed via reverse-phase UHPLC as described in Weigert *et al*.^40^

### *p*NPB-activity measurement for thermostability assessment of *Halo*PETase1

500 nM of *Halo*PETase1 were incubated in 1 ml 20 mM NaPi, pH 7.4, with 1, 3 or 5 M NaCl concentration at 50, 60 or 68 °C for up to 72 h. 10 µl samples were taken after 1, 4, 24, 48, and 72 h from each salt- and temperature-condition and diluted with 980 µl 20 mM NaPi pH 7.4, 150 mM NaCl in a UV-transmissible plastic cuvette. To begin the hydrolytic reaction, 10 µl of a 100 mM pNPB in DMSO were transferred to the enzyme solution in the cuvette. The reaction was mixed by inversion and kinetic absorbance measurement were performed at 405 nm immediately after mixing. The residual activity was determined by normalizing the initial rates with the one at 0 h incubation.

### Heterologous expression and purification of *Halo*PETases2 to 6

The genes of *Halo*PETases2 to 6 were analyzed with SignalP 6.0^49^ to identify and remove the N-terminal signal peptide. Genes were codon-optimized for production in *E. coli* Shuffle^®^ T7 cells (NEB) and cloned into the pET21b-vector. Pre-cultures were prepared as before. The expression was then induced by addition of 0.1 mM IPTG (final concentration) and cultures were left overnight at 16 or 20 °C with 140 rpm shaking. Cells were harvested and pellets washed with 30 ml PBS.

Cells were thawed on ice, resuspended with 30 ml of buffer A and subsequently lysed via ultrasonication. Crude cell lysates were centrifuged to separate cell debris from soluble protein at 4 °C, 18000 rpm for 1 h. The cleared lysates (supernatants) were transferred into a fresh vessel and stored on ice. Ni-NTA and SEC purification steps were performed for all proteins. If insufficient purity was observed by SDS-PAGE, ion-exchange chromatography was performed as a final step.

If so, fractions from the SEC were pooled and concentrated to dilute the NaCl concentration 20-fold from 150 to theoretically 7.5 mM with buffer D (20 mM NaPi, pH 7.4). Cation- and anion-exchange chromatography were performed with a 1 ml HiTrap^®^ SP FF or 1 ml HiTrap^®^ Q FF column, respectively. The elution was performed by a linear gradient from 0 to 25 % of buffer E (20 mM NaPi, 2 M NaCl, pH 7.4) over 20 CV. The peak fractions were pooled, concentrated, aliquoted and flash frozen.

### Site-directed mutagenesis (SDM) of *Halo*PETase1

Single mutations were introduced in *Halo*PETase1 at position T88 to re-establish the π-stacking clamp and at position Q157 to re-introduce the canonical methionine in the lipase-box motif using site-directed mutagenesis. See Table S5 for primer pairs and sequences. Sequences were verified by Sanger sequencing.

### Bioinformatic mining for putative PETases in *Halopseudomonas*

A search was conducted against the proteome of *Halopseudomonas*. For this, 40 genomes were retrieved from NCBI (for accession numbers see Table S4). Open reading frames of the *Halopseudomonas* genomes were predicted with Prodigal version 2.6.3^53^. A hidden Markov model profile from biochemically characterized PETases from the PAZy database^29^ was generated with HMMER version 3.1b1^48^ using the *hmmbuild* option and a search against the *Halopseudomonas* proteome was performed using HMMER 3.1b1 with the *hmmsearch* option and a bitscore threshold of 100.

### Three-dimensional structure determination

*Halo*PETase1 in 50 mM Tris, 150 mM NaCl, pH 7.5 was set to a final concentration of 10 mg/ml. Sitting-drop vapor-diffusion methodology was used with JCSG Core I-III and PEGs II (Qiagen) for screening crystallization conditions in 96 well Intelli plates (Art Robbins) stored at 20 °C in Rock Imager RI 182 (Formulatrix). Volume ratios of 2:1, 1:1, and 1:2 were prepared using a Phoenix robot dispenser (Art Robbins Instruments) with a total drop volume of 0.8 µl. Crystal formation was observed in the following condition: 1:1 ratio, 0.1 M imidazole, 40 % (v/v) PEG 400. For cryoprotection 25 % (v/v) glycerol was added to the crystals before flash-freezing in liquid nitrogen. Diffraction data were collected at the German Electron Synchroton (DESY) beamline P13^54^ operated by the Helmholtz association using a wavelength of 0.97 Å and a PILATUS detector.

Diffraction data were processed with X-ray Detector Software (XDS) using XDSAPP 3.1.5^55^. The structure was solved by molecular replacement with PHASER in the PHENIX^56^ software suite v.1.21 using a ColabFold^45^ model of *Halo*PETase1. The refinement was performed with phenix.refine. The model was improved by electron density map inspection and iterative manual rebuilding performed in Coot v.0.9^57^. The 3D-structure and structure factors were deposited in the PDB^58^ with accession code 9hl5. All structural representations and graphical manipulations were performed in PyMOL 3.0.0 (Schrodinger, LLC)^59^.

### Sequence and structure analysis of *Halo*PETases

Multiple sequence alignments (MSAs) were performed using ProbCons^60^ with full-length sequences. Alignment of *Pseudomonas mendocina* cutinase (*Pm*C) was performed with the protein sequence retrieved from UniProtKB^61^ (entry: A4Y035). Sequence homology analysis of *Halo*PETases and reference PETases such as leaf-branch compost cutinase (LCC, UniProtKB entry: G9BY57) and *Ideonella sakaiensis* PETase (*Is*PETase, UniProtKB entry: A0K8P6T7) was performed by creating a MSA with ProbCons and using the ClustalX 2.1^62^ software suite to generate a neighborhood-join tree with 1000 bootstrap. The tree was illustrated using the website tool Interactive Tree of Life (iTOL, https://itol.embl.de/). To plot the amino-acid frequencies from the MSA as a logo plot, the python library LogoMaker was employed (https://github.com/jbkinney/logomaker).

The 3D structures of putative and experimentally characterized *Halo*PETases were modeled with ColabFold^45^, only rank 1 structures were considered for further analyses. Structural alignments were generated in PyMOL with the *align* algorithm (default settings; 2 Å cut-off and 5 refinement cycles) implemented in the software. In all cases, the alignment was done globally and with all backbone C_α_ atoms, except the first 40 C_α_ to exclude the N-terminal signal peptides.

### Preparation of PET-films

Polymer films were prepared using a 25-12-2HC hot press (Carver) with a 130x130x0.1 mm mold. The required amount of polymer (fibre grade PET granules RT3050, Indorma Ventures) was placed in the mold, which was then sandwiched between stainless-steel plates coated with Nowoflon Perfluoralkoxy (PFA) foil. The material was heated to 270°C for 3 minutes and pressed at 5 tons for 1 minute. Afterwards, the PET films, along with the mold, were rapidly quenched in ice water immediately after removal from the hot press. The crystallinity of the films was determined using DSC.

### Differential scanning calorimetry (DSC)

Differential scanning calorimetry (DSC) measurements were conducted using a Netzsch 204 F1 Phoenix instrument. Approximately 5-7 mg of the sample was placed in an aluminum crucible with a pierced lid. The analysis was performed over a temperature range of 0 to 300°C under a nitrogen atmosphere (Flow: 20 ml/min). Data analysis was carried out using the Proteus Analysis 8.0 software.

The percentage crystallinity of the polymer was determined using DSC by calculating the ratio of the enthalpy of fusion of the polymer sample subtracted by enthalpy of crystallization to the enthalpy of fusion of the polymer at 100% crystallinity, as shown in the following equation:

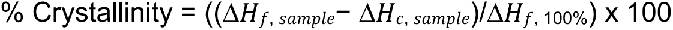

Where Δ*H*_*f, sample*_, is the enthalpy of fusion of the sample, Δ*H*_*c, sample*_ is the enthalpy of crystallization obtained from the 1^st^ heating curve of the DSC plot. Δ*H*_*f*, 100%_ represents the enthalpy of fusion of the fully crystalline polymer. The Δ*H*_*f*, 100%_ was taken as 140 J/g^63^. Using the above equation the crystallinity of the PET films was calculated to be ∼14%.

### Scanning electron microscopy (SEM)

SEM was performed on an FEI Quanta FEG 250 (Thermo Fisher Scientific), equipped with an ET detector. To prepare the samples for imaging, a 2.0 nm platinum coating was applied using a Cressington 208HR Platinum Sputter Coater, with the coating thickness monitored by an MTM20 thickness monitor.

---

**Figure S1:**
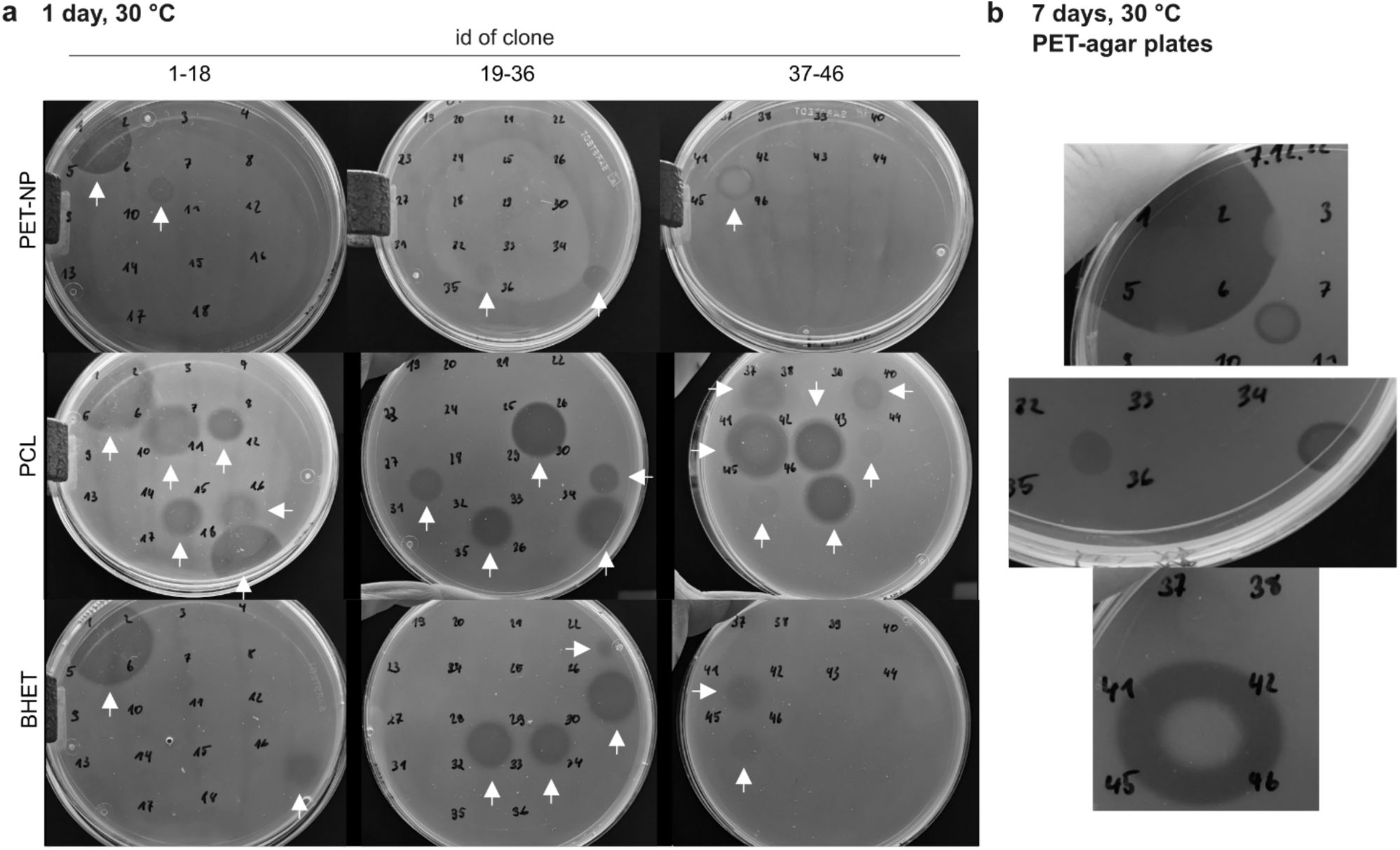
Functional screening of target proteins via plate-based assay. PBS-agar (1.5 % [w/v] agar) plates were supplemented with either PET-NP, PCL or BHET and purified enzyme solutions were added on the agar surface. **a)** Zone of clearances on all plates were observed after 1 day **b)** and on PET-NP after 7 days of incubation at 30 °C. IDs next to the spots and above in **a** and **b** represent an individual recombinant target protein as shown in Table S1.

**Figure S2:**
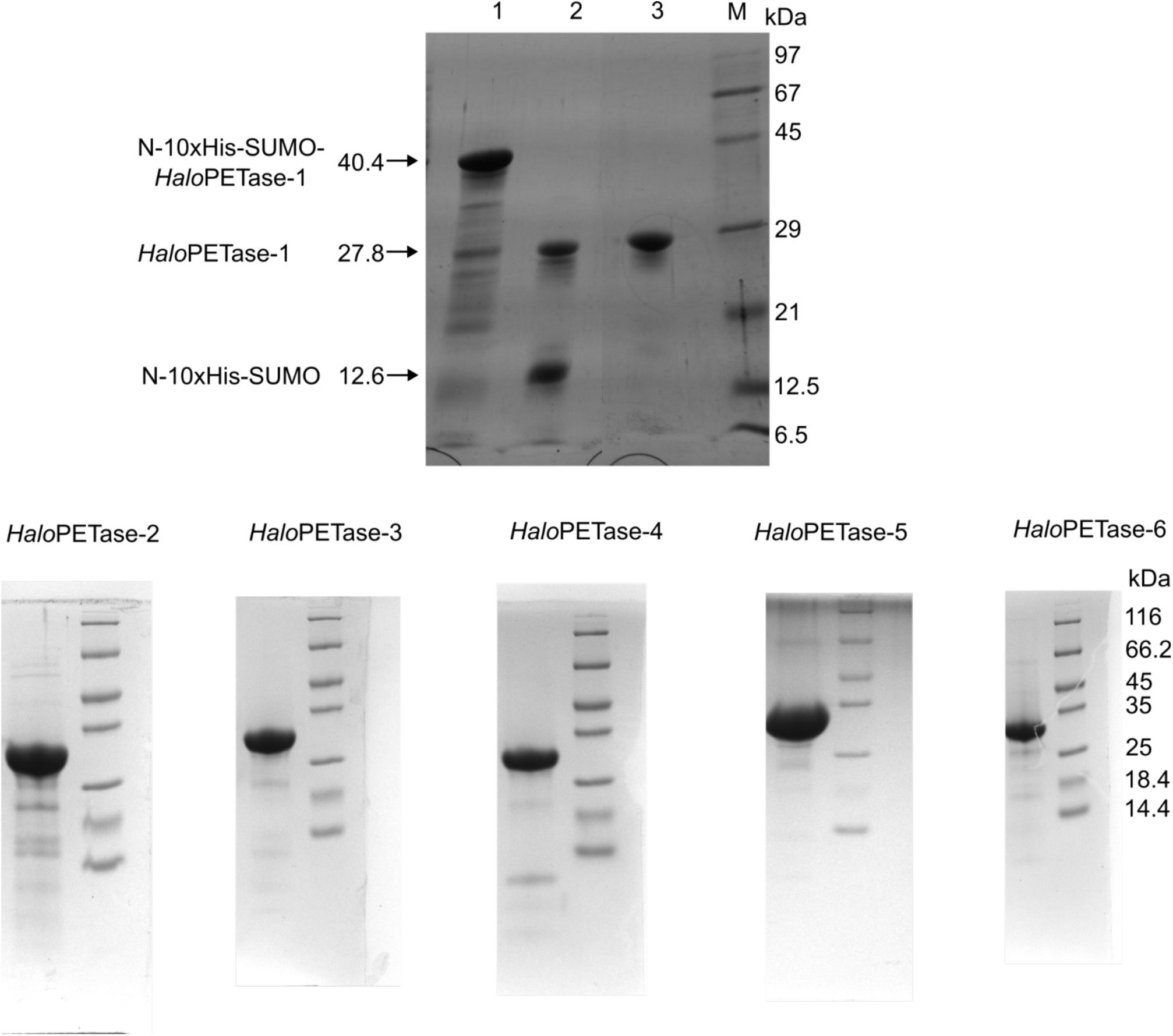
SDS-PAGE gels of *Halo*PETase purifications. Upper gel: Purification of N-10xHis- SUMO-tagged *Halo*PETase-1 (PSW62-2) by Ni-IMAC (lane 1), SenP2-digestion and dialysis for the removal of N-10xHis-SUMO-tag (lane 2), reverse Ni-IMAC and SEC (lane 3) to obtain truncated and purified *Halo*PETase1. Lower gels: Final purified protein solutions of *Halo*PETases2 to 6.

**Figure S3:**
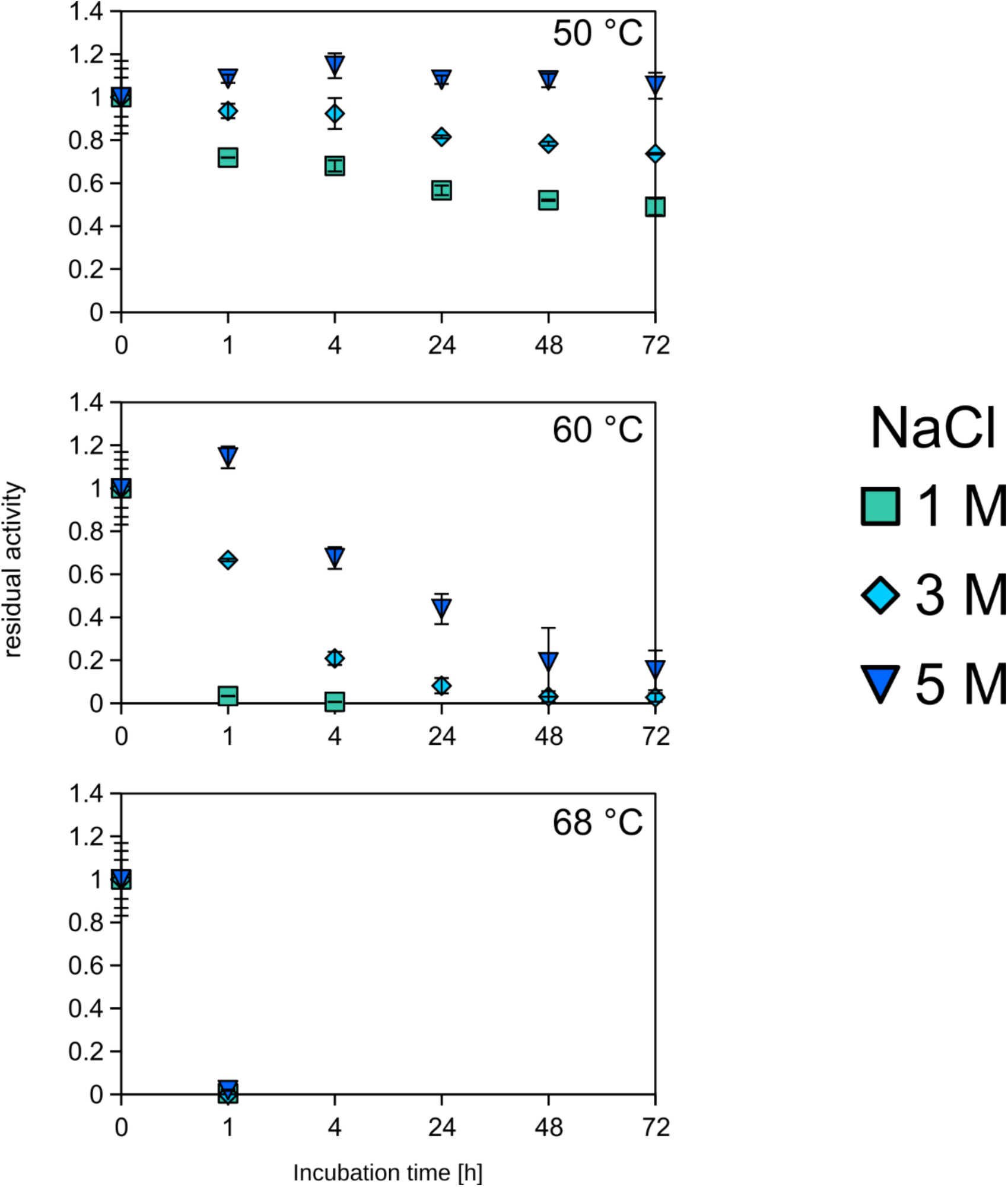
Residual activity of *Halo*PETase1 tested via *p*NPB hydrolysis depending on NaCl concentration, time and temperature. The residual activity was assessed by measuring the absorbance of the hydrolysis product *p*NP at 405 nm after incubation under specific salt and temperature conditions and normalizing the change at A405nm over time to the initial rate at 0 hours. Enzyme solutions were prepared with 5 µM concentration in the respective condition and incubated for up to 72 h. Subsequently, incubated solutions were diluted 100-fold in reaction solution, containing 20 mM sodium phosphate, 150 mM NaCl with pH 7.4 and 1 mM *p*NPB.

**Figure S4:**
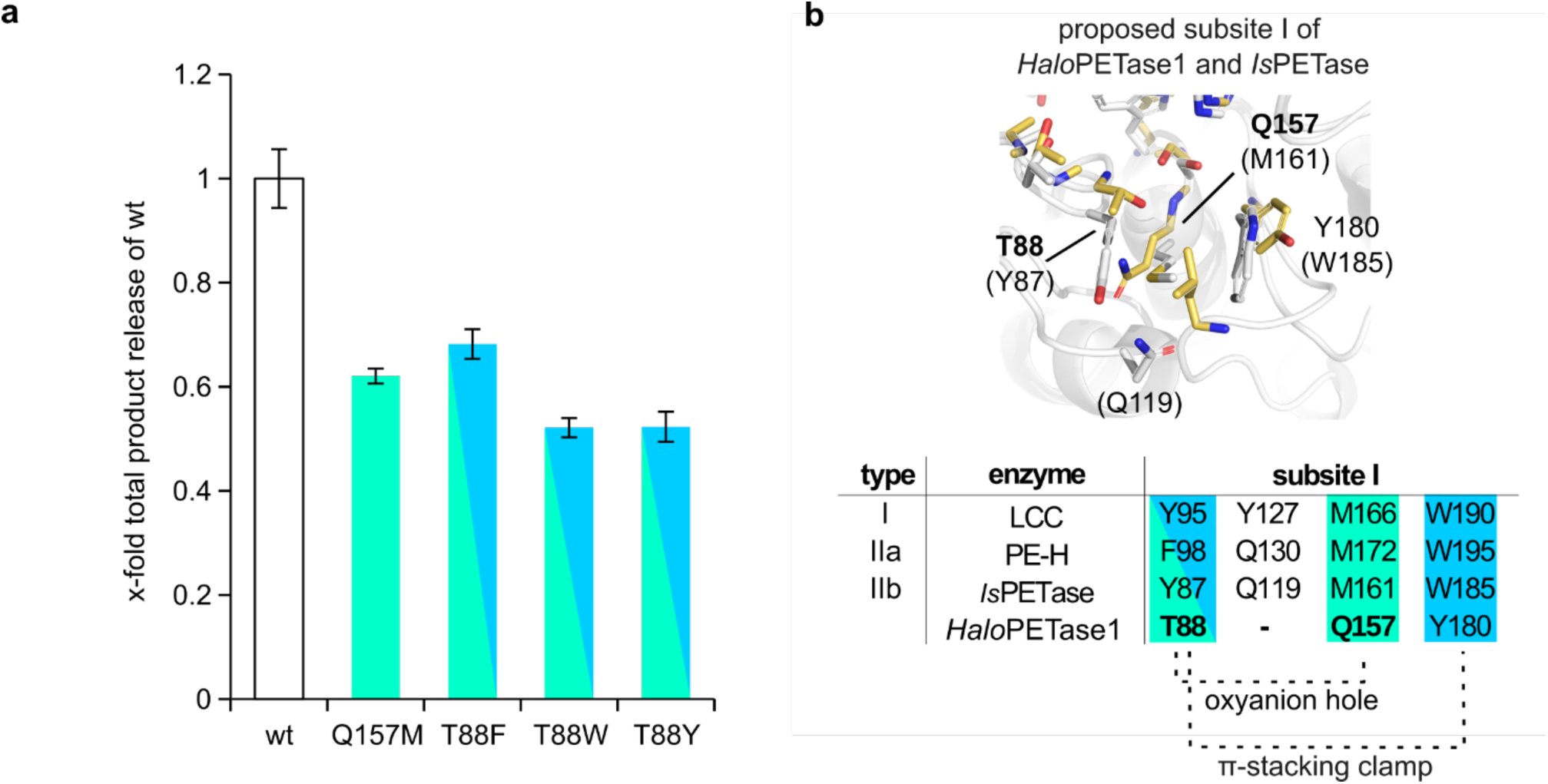
Effect of single mutations in the subsite I of *Halo*PETase1. **a**) PET-degradation by *Halo*PETase1 (wt) and its variants in a PET-coated well-plate assay. T88 was substituted with an aromatic amino acid (T88F, T88W, T88Y) to restore the canonical π-stacking clamp. Q157 was substituted by methionine (Q157M) to obtain the representative occupation in the oxyanion hole, as in other PETases. Reaction buffer composition: 20 mM NaPi, 3 M NaCl, pH 7.4. The reactions were incubated at 50 °C for 20 h. **b**) Aligned subsite I of *Halo*PETase1 and *Is*PETase. Corresponding amino acids highlighted as stick representations (yellow: *Halo*PETase1, grey: *Is*PETase). Subsite I amino acid occupation from type I, IIa, IIb and *Halo*PETase1 is shown in the table, where significant substitutions in *Halo*PETase1 are highlighted with bold letters. L163 of *Halo*PETase1 in the vicinity of Q157 and Y180 was not categorized and thus not labelled.

**Figure S5:**
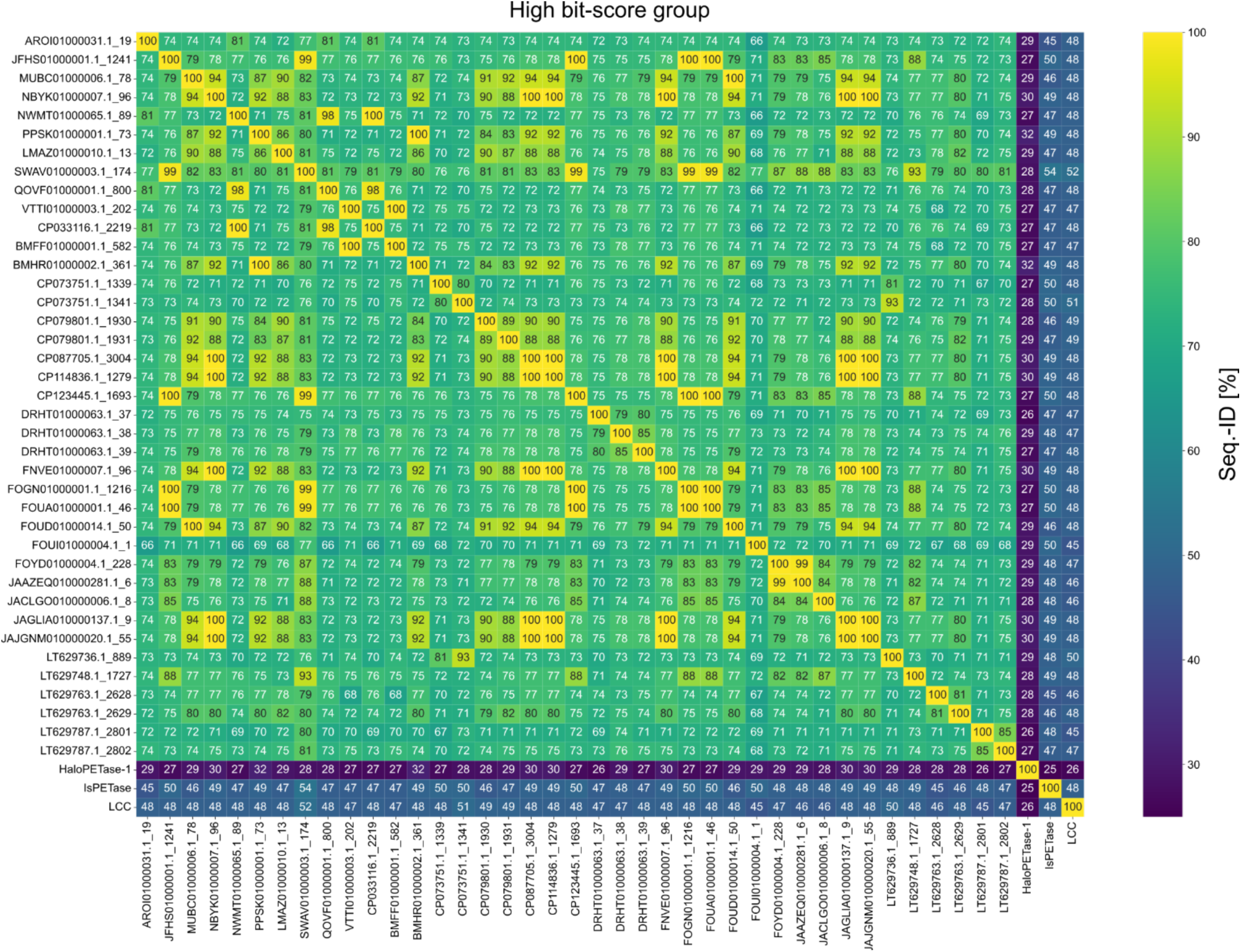
Sequence identity matrix of pairwise alignments with candidate PETases from the high bit-score group. All aligned sequences were included in the phylogenetic tree in Figure 3a. Pairwise alignments were performed by ProbCons and sequence identities were inferred from ClustalX^5^.

**Figure S6:**
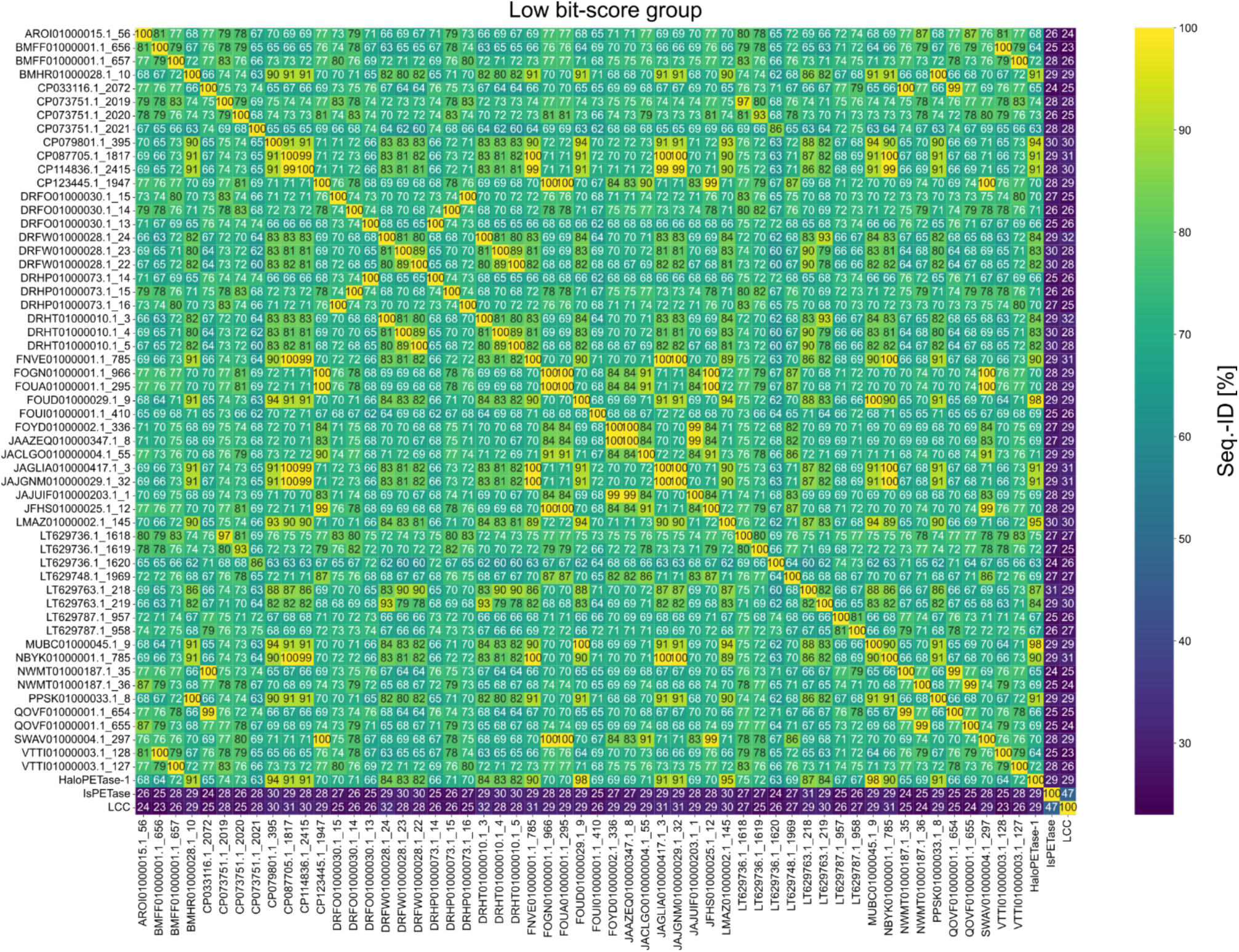
Sequence identity matrix of pairwise alignments with candidate PETases from the low bit-score group. All aligned sequences were included in the phylogenetic tree in Figure 3a. Pairwise alignments were performed by ProbCons and sequence identities were inferred from ClustalX^5^.

**Figure S7:**
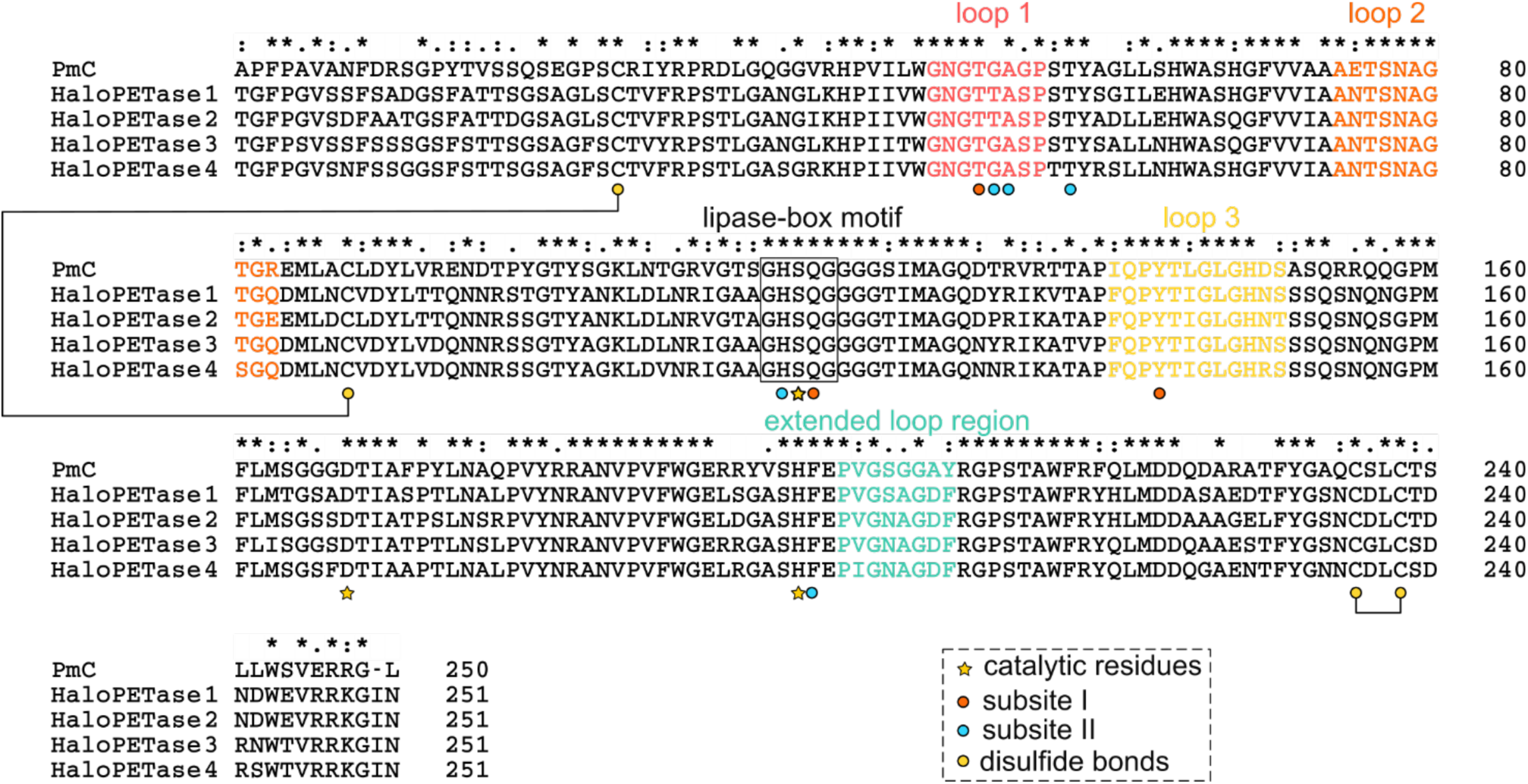
MSA of low bit-score PETases including *Halo*PETase1 to 4 and *Pm*C. Numbers at the end of each line represent the sequence position of the last amino acid of the corresponding protein from each line. The extent of conservation at each position is indicated by ‘*’ for identical, ‘:’ for highly similar or ‘.’ less similar residues, according to the Gonnet PAM 250 scoring matrix^6^. Colored circle or star symbols indicate potentially important and catalytic residues at the active site, respectively. Connected yellow circles represent cysteines forming a disulfide bond. Disulfide bonds from *Halo*PETase2 to 4 are inferred from *Halo*PETase1. The extended loop region (turquoise-colored letters) was inferred from Joo *et al*.^7^. Loops characterized in this work (loop 1 to 3) are colored according to Figure 3.

**Figure S8:**
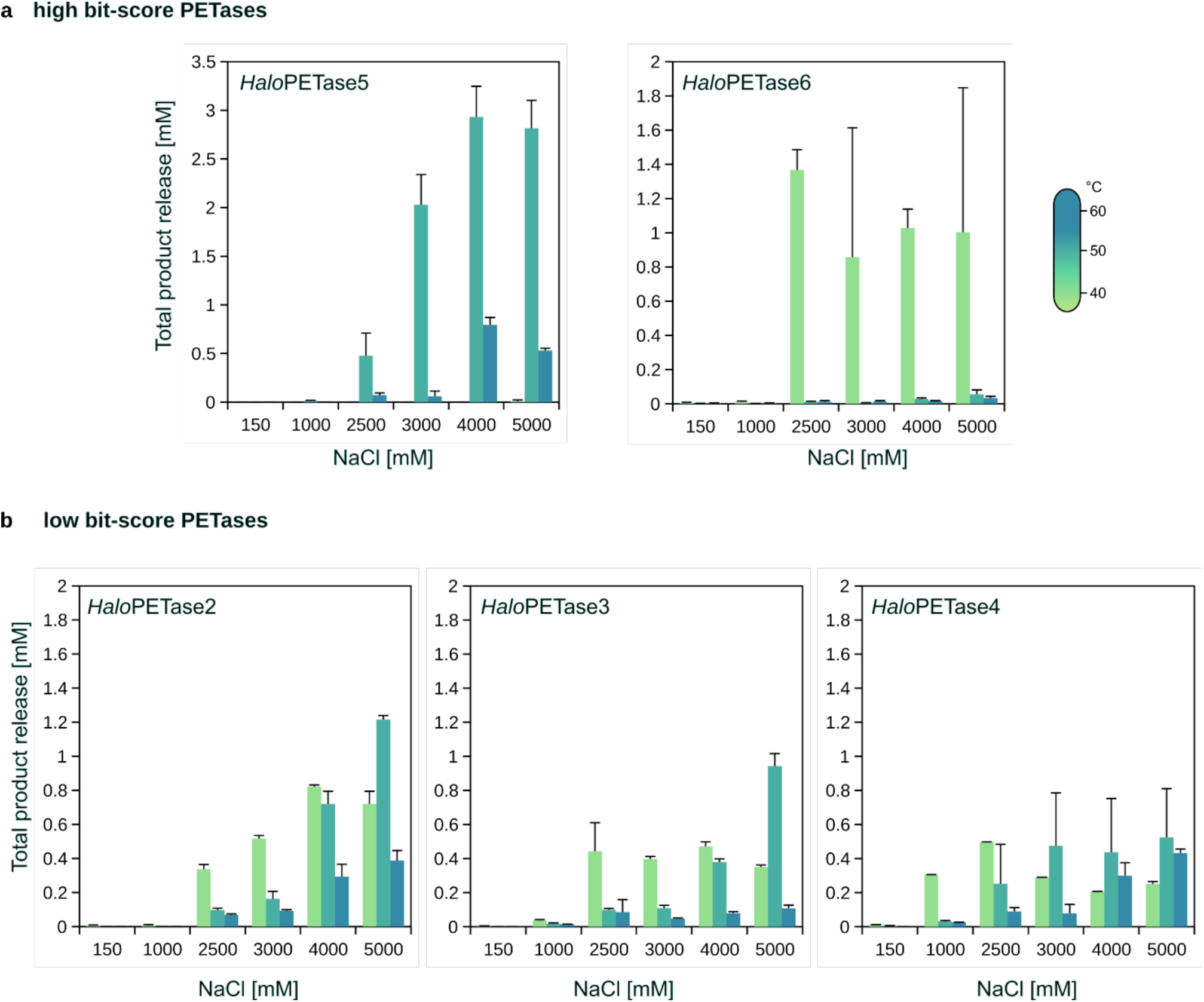
Salt- and temperature-dependent PET-degradation by *Halo*PETase2 to 6 from low and high bit-score groups. Degradation experiments were performed with the PET-coated assay according to Weigert *et al.* (see Methods section). The buffer for each experiment was 20 mM NaPi, pH 7.4 with variable NaCl-concentrations. Protein concentrations were selected according to preliminary testing for optimal concentration as follows: **a)** high bit-score *Halo*PETases: *Halo*PETase5 550 nM; *Halo*PETase6: 50 nM. **b)** low bit-score *Halo*PETases: *Halo*PETase2 500 nM; *Halo*PETase3 550 nM; *Halo*PETase4 150 nM. Error-bars indicate standard deviations of triplicate measurements.

**Figure S9:**
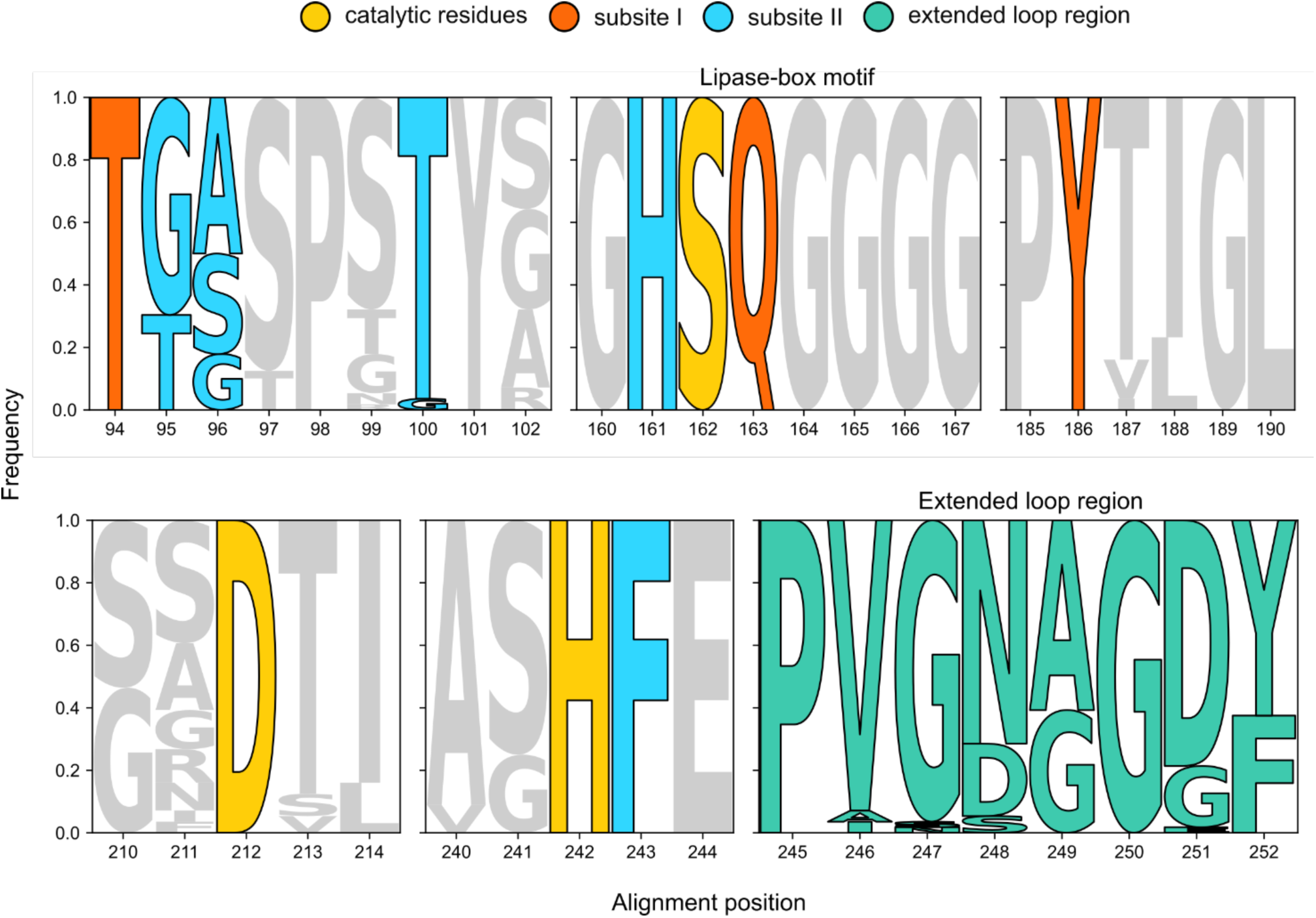
Logo plot of important regions in *Halo*PETases from the lower bit-score group. Important residues and regions for PETase activity according to Joo *et al*.^7^. The logo-plot was generated using full-length sequences of putative low bit-score PETase sequences from *Halopseudomonas*, as listed in Table S4.

**Figure S10:**
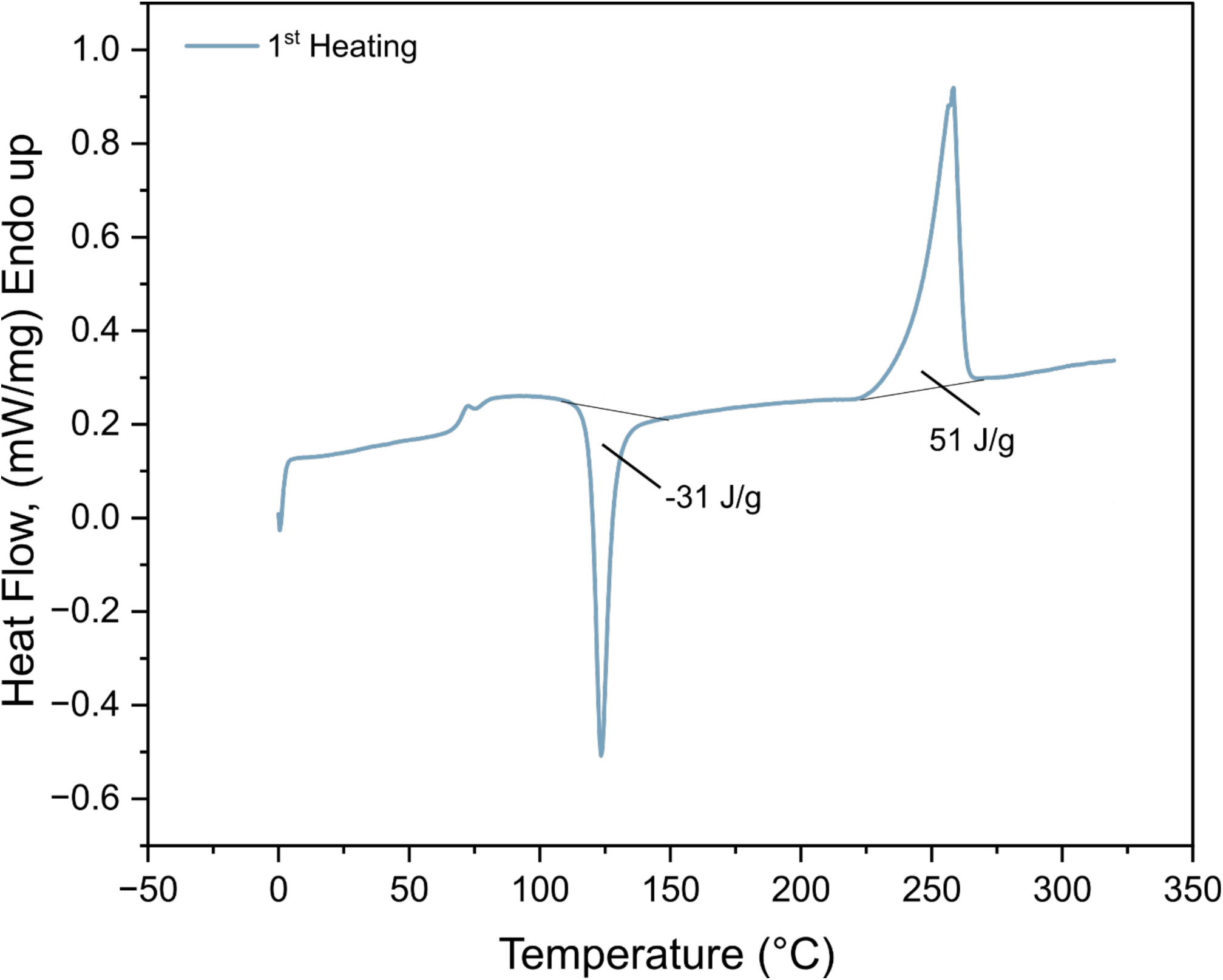
PET-film characterization by DSC. Calculation of crystallinity was performed using the first heating curve. More specifically, the heat of crystallization (31 J/g) was subtracted from the heat of fusion (45 J/g) and divided by the heat of fusion for 100 % crystalline PET (see Methods).

**Table S1:** Functional plate-based screening of recombinant target proteins. The purified protein IDs correspond to the tested proteins in Figure S1. Target protein ID 1 corresponds to the positive control, IDs 2-5 to LCC(ICCG) and negative controls, i.e. GSGSGS peptide sequence attached to one of the possible purification and/or solubility tags. A ‘+’ represents qualitatively detected zone of clearance and ‘-’ no detection of the same. PSW_62_2 highlighted in bold) is the screening name for *Halo*PETase1.

| Purified protein ID | Target protein | Tag | PCL | BHET | PET-NP |
| --- | --- | --- | --- | --- | --- |
| 1 | LCC(ICCG) | C-6xHis | + | + | + |
| 2 | GSGSGS | N-10xHis | - | - | - |
| 3 | GSGSGS | C-6xHis | - | - | - |
| 4 | GSGSGS | N-10xHis-MBP | - | - | - |
| 5 | GSGSGS | N-10xHis-SUMO | - | - | - |
| 6 | **PSW_62_2** | N-10xHis | + | - | + |
| 7 | ANW_6_2 | N-10xHis | + | - | - |
| 8 | PSW_64_1 | N-10xHis | - | - | - |
| 9 | ION_47_1 | N-10xHis | - | - | - |
| 10 | ASW_18_1 | N-10xHis | + | - | - |
| 11 | IOS_14_1 | N-10xHis | + | - | - |
| 12 | PSE_95_1 | N-10xHis | - | - | - |
| 13 | P01_A02_3_1 | N-10xHis | - | - | - |
| 14 | PSE_112_1 | N-10xHis | + | - | - |
| 15 | ANW_30_1 | N-10xHis | + | - | - |
| 16 | PSE_138_1 | C-6xHis | + | + | - |
| 17 | **PSW_62_2** | C-6xHis | - | - | - |
| 18 | ANW_6_2 | C-6xHis | + | - | - |
| 19 | IOS_57_1 | C-6xHis | - | - | - |
| 20 | ION_47_1 | C-6xHis | - | - | - |
| 21 | ASW_18_1 | C-6xHis | - | - | - |
| 22 | IOS_14_1 | C-6xHis | - | + | - |
| 23 | PSE_95_1 | C-6xHis | - | - | - |
| 24 | P01_A02_3_1 | C-6xHis | - | - | - |
| 25 | PSE_112_1 | C-6xHis | + | - | - |
| 26 | SWF_bin8_13 | C-6xHis | - | + | - |
| 27 | ANW_30_1 | C-6xHis | + | - | - |
| 28 | IOS_26_1 | C-6xHis | - | + | - |
| 29 | PON_52_1 | C-6xHis | - | + | - |
| 30 | PSE_138_1 | N-10xHis-MBP | - | + | - |
| 31 | PSE_120_1 | N-10xHis | - | - | - |
| 32 | PSW_62_1 | N-10xHis | + | - | + |
| 33 | PSE_130_1 | N-10xHis | + | - | - |
| 34 | ASW_29_1 | N-10xHis | + | - | + |
| 35 | ANW_6_1 | N-10xHis | - | - | - |
| 36 | PSE_120_1 | C-6xHis | + | - | - |
| 37 | PSW_62_1 | C-6xHis | + | - | + |
| 38 | PSE_130_1 | C-6xHis | - | - | - |
| 39 | ASW_29_1 | C-6xHis | + | - | - |
| 40 | ANW_6_1 | C-6xHis | - | - | - |
| 41 | **PSW_62_2** | N-10xHis-SUMO | + | + | + |
| 42 | ANW_6_2 | N-10xHis-SUMO | + | - | - |
| 43 | ASW_18_1 | N-10xHis-SUMO | + | - | - |
| 44 | PSE_95_1 | N-10xHis-SUMO | - | - | - |
| 45 | P01_A02_3_1 | N-10xHis-SUMO | + | + | - |
| 46 | PSE_112_1 | N-10xHis-SUMO | + | - | - |

**Table S2:**
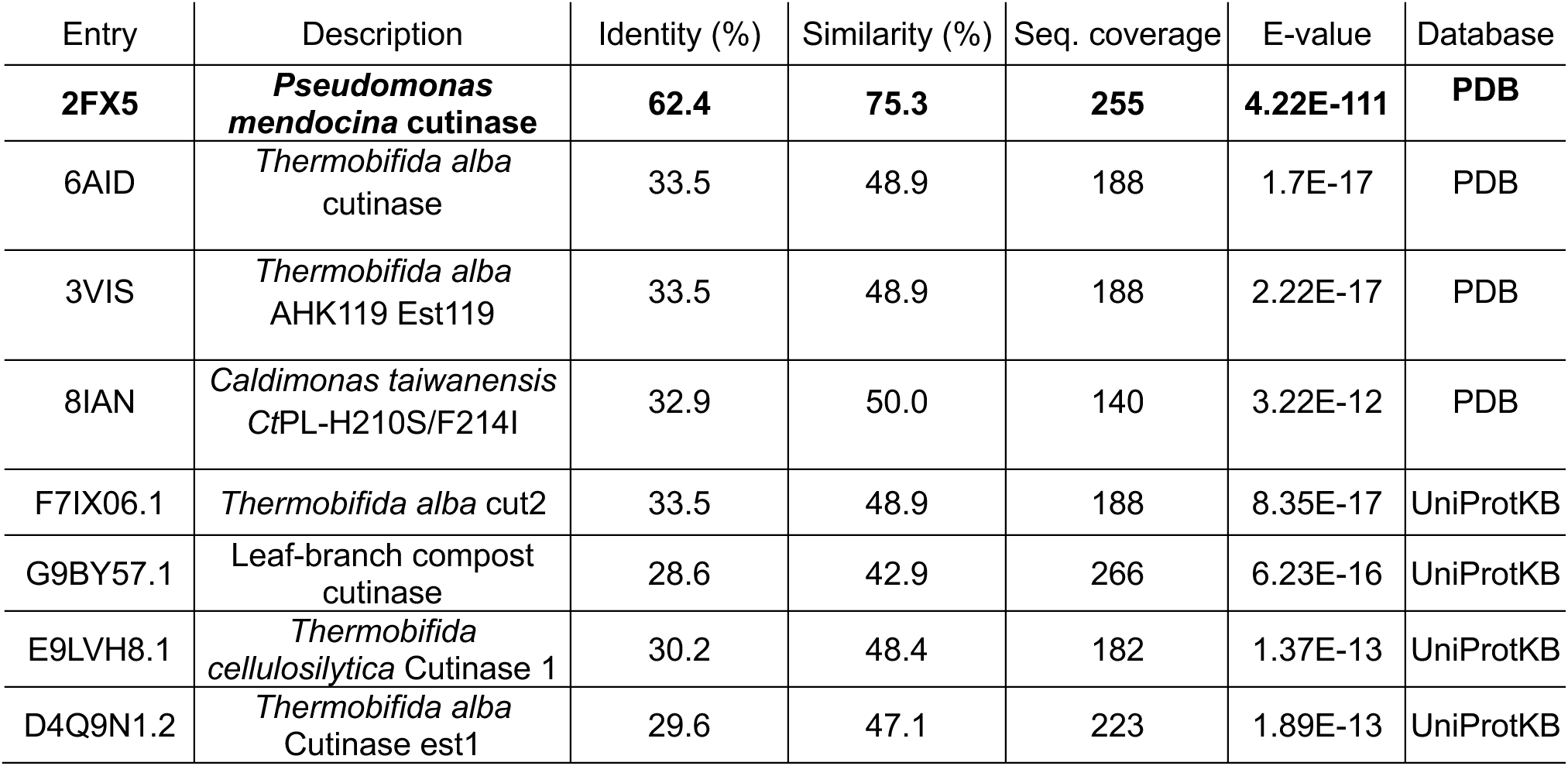
BLASTp hits for *Halo*PETase1 in the databases PDB and UniProtKB. The analysis was performed on August 6^th^, 2024. The hit with highest sequence identity, *Pseudomonas mendocina c*utinase, is highlighted in bold.

| Entry | Description | Identity (%) | Similarity (%) | Seq. coverage | E-value | Database |
| --- | --- | --- | --- | --- | --- | --- |
| **2FX5** | ***Pseudomonas mendocina* cutinase** | **62.4** | **75.3** | **255** | **4.22E-111** | **PDB** |
| 6AID | *Thermobifida alba* cutinase | 33.5 | 48.9 | 188 | 1.7E-17 | PDB |
| 3VIS | *Thermobifida alba* AHK119 Est119 | 33.5 | 48.9 | 188 | 2.22E-17 | PDB |
| 8IAN | *Caldimonas taiwanensis* CtPL-H210S/F214I | 32.9 | 50.0 | 140 | 3.22E-12 | PDB |
| F7IX06.1 | *Thermobifida alba* cut2 | 33.5 | 48.9 | 188 | 8.35E-17 | UniProtKB |
| G9BY57.1 | Leaf-branch compost cutinase | 28.6 | 42.9 | 266 | 6.23E-16 | UniProtKB |
| E9LVH8.1 | *Thermobifida cellulosilytica* Cutinase 1 | 30.2 | 48.4 | 182 | 1.37E-13 | UniProtKB |
| D4Q9N1.2 | *Thermobifida alba* Cutinase est1 | 29.6 | 47.1 | 223 | 1.89E-13 | UniProtKB |

**Table S3:**
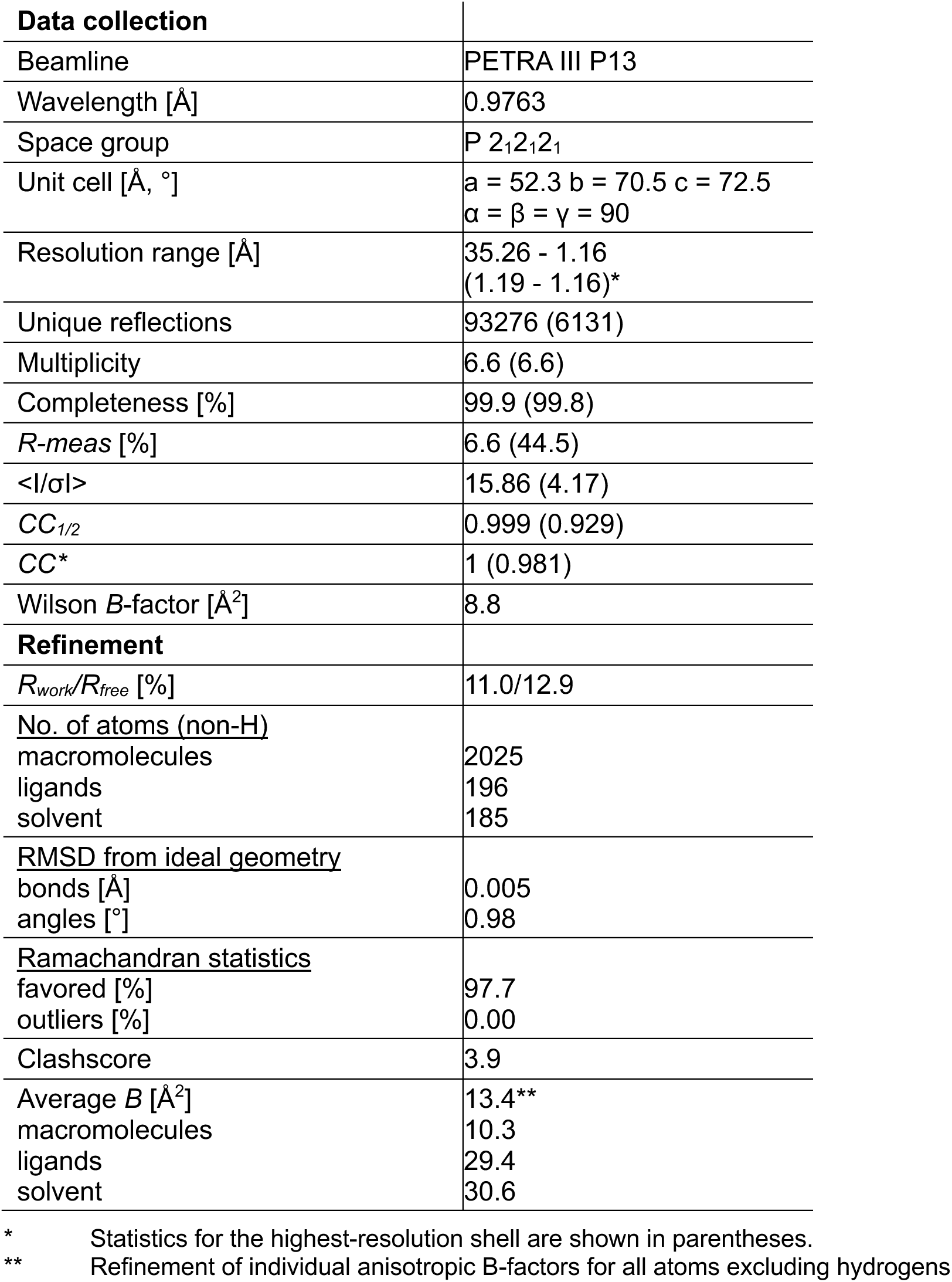
Crystallographic data collection and refinement statistics.

**Table S4:** Results of *Halo*PETase1 homolog mining in *Halopseudomonas* using pHMM analysis. The pHMM analysis was performed in *Halopseudomonas* genome assemblies, which can be found by their NCBI Genome GenBank^1^ accession IDs. Corresponding hits are shown in the **‘**Protein designation’ column, whereby the IDs correspond to the position of the protein on the specific contig. Hit sequences were affiliated with the low bit-score group if the respective bit-score was lower than 200. Otherwise, hits belong to the high bit-score group. Length in number of amino acids in one hit sequence corresponds to a full sequence including possible N-terminal signal peptide sequence. Biochemically characterized hits are referred to a publication or to this work in the column ‘Characterized’. Characterized proteins were verified through a BLASTp^2^ search against non-redundant protein databases on September 11, 2024.

| GenBank genome accession | Protein designation | Affiliation | Length (aa) | Bit-score | Bit-score group | PETase type | Characterized |
| --- | --- | --- | --- | --- | --- | --- | --- |
| GCA_020781875.1 | JAJGNM010000029.1 32 | H. aestusnigri FXH-240 | 290 | 143.8 | Low | Putative III | - |
| GCA_020781875.1 | JAJGNM010000020.1 55 | H. aestusnigri FXH-240 | 304 | 395.2 | High | Putative IIa | - |
| GCA_021184005.1 | CP087705.1 1817 | H. aestusnigri GOM5 | 290 | 143.8 | Low | Putative III | - |
| GCA_021184005.1 | CP087705.1 3004 | H. aestusnigri GOM5 | 304 | 395.2 | High | Putative IIa | - |
| GCA_023143355.1 | JAGLIA010000417.1 3 | H. aestusnigri RS 2 23 | 290 | 143.8 | Low | Putative III | - |
| GCA_023143355.1 | JAGLIA010000137.1 9 | H. aestusnigri RS 2 23 | 304 | 395.2 | High | Putative IIa | - |
| GCA_002197985.1 | NBYK01000001.1 785 | H. aestusnigri VGXO14 | 289 | 143.7 | Low | Putative III | - |
| GCA_002197985.1 | NBYK01000007.1 96 | H. aestusnigri VGXO14 | 304 | 396.3 | High | IIa | *Haes*_PE-H<sup>3</sup> |
| GCA_005096325.1 | SWAV01000004.1 297 | H. bauzanensis SBBB | 287 | 140 | Low | Putative III | - |
| GCA_005096325.1 | SWAV01000003.1 174 | H. bauzanensis SBBB | 196 | 288.2 | High | IIa | *Pbauz*Cut<sup>4</sup> |
| GCA_000632535.1 | JFHS01000025.1 12 | H. bauzanensis W13Z2 | 287 | 142.8 | Low | Putative III | - |
| GCA_000632535.1 | JFHS01000001.1 1241 | H. bauzanensis W13Z2 | 302 | 398.9 | High | IIa | *Pbauz*Cut<sup>4</sup> |
| GCA_012512195.1 | JAAZEQ010000347.1 8 | H. formosensis AS09scLD 420 | 287 | 143.8 | Low | Putative III | - |
| GCA_012512195.1 | JAAZEQ010000281.1 6 | H. formosensis AS09scLD 420 | 304 | 392.3 | High | Putative IIa | - |
| GCA_029263535.1 | JAJUIF010000203.1 1 | H. formosensis bin83 | 287 | 144.6 | Low | Putative III | - |
| GCA_029263535.1 | JAJUIF010000365.1 5 | H. formosensis bin83 | 129 | 162.2 | Low | Putative III | - |
| GCA_003444685.1 | LMAZ01000002.1 145 | H. gallaeciensis V113 | 289 | 139.3 | Low | Putative III | - |
| GCA_003444685.1 | LMAZ01000010.1 13 | H. gallaeciensis V113 | 305 | 383.3 | High | Putative IIa | - |
| GCA_008365385.1 | QOVF01000001.1 655 | H. laoshanensis Y22 | 288 | 139.8 | Low | Putative III | - |
| GCA_008365385.1 | QOVF01000001.1 654 | H. laoshanensis Y22 | 287 | 146.3 | Low | Putative III | - |
| GCA_008365385.1 | QOVF01000001.1 800 | H. laoshanensis Y22 | 310 | 402.2 | High | Putative IIa | - |
| GCA_002903165.1 | PPSK01000033.1 8 | H. oceani DSM 100277 | 288 | 132.8 | Low | Putative III | - |
| GCA_002903165.1 | PPSK01000001.1 73 | H. oceani DSM 100277 | 303 | 395.2 | High | Putative IIa | - |
| GCA_001989375.1 | MUBC01000045.1 9 | H. pachastrellae CCUG 46540 | 289 | 139.6 | Low | Putative III | - |
| GCA_001989375.1 | MUBC01000006.1 78 | H. pachastrellae CCUG 46540 | 304 | 387 | High | Putative IIa | - |
| GCA_015661625.1 | DQVS01000080.1 1 | H. pachastrellae SZUA-1527 | 127 | 154.5 | Low | Putative III | - |
| GCA_002351465.1 | NWMT01000187.1 36 | H. pelagia 58 153 | 288 | 139.2 | Low | Putative III | - |
| GCA_002351465.1 | NWMT01000187.1 35 | H. pelagia 58 153 | 286 | 147.4 | Low | Putative III | - |
| GCA_002351465.1 | NWMT01000065.1 89 | H. pelagia 58 153 | 310 | 404.3 | High | Putative IIa | - |
| GCA_000410875.1 | AROIO1000015.1 56 | H. pelagia CL-AP6 | 285 | 139.6 | Low | Putative III | - |
| GCA_000410875.1 | AROIO1000031.1 19 | H. pelagia CL-AP6 | 307 | 412.1 | High | Putative IIa | - |
| GCA_009497895.1 | CP033116.1 2073 | H. pelagia Kongs-67 | 288 | 139.2 | Low | Putative III | - |
| GCA_009497895.1 | CP033116.1 2072 | H. pelagia Kongs-67 | 286 | 147.4 | Low | Putative III | - |
| GCA_009497895.1 | CP033116.1 2219 | H. pelagia Kongs-67 | 310 | 404.3 | High | Putative IIa | - |
| GCA_011050335.1 | DRFW01000028.1 22 | H. sabulinigri HyVt-331 | 288 | 139.4 | Low | Putative III | - |
| GCA_011050335.1 | DRFW01000028.1 23 | H. sabulinigri HyVt-331 | 290 | 140.6 | Low | Putative III | - |
| GCA_011050335.1 | DRFW01000028.1 24 | H. sabulinigri HyVt-331 | 289 | 148.2 | Low | Putative III | - |
| **GCA_011053055.1** | **DRHT01000010.1 5** | **H. sabulinigri HyVt-376** | **288** | **139.4** | **Low** | III | **This work** |
| **GCA_011053055.1** | **DRHT01000010.1 4** | **H. sabulinigri HyVt-376** | **290** | **140.6** | **Low** | III | **This work** |
| **GCA_011053055.1** | **DRHT01000010.1 3** | **H. sabulinigri HyVt-376** | **289** | **148.2** | **Low** | III | **This work** |
| GCA_011053055.1 | DRHT01000063.1 37 | H. sabulinigri HyVt-376 | 300 | 385.5 | High | Putative IIa | - |
| **GCA_011053055.1** | **DRHT01000063.1 39** | **H. sabulinigri HyVt-376** | **301** | **388.9** | **High** | **Putative IIa** | **This work** |
| **GCA_011053055.1** | **DRHT01000063.1 38** | **H. sabulinigri HyVt-376** | **305** | **391.2** | **High** | **Putative IIa** | **This work** |
| GCA_008641105.1 | VTTIO1000003.1 127 | H. salina XCD-X85 | 293 | 134.2 | Low | Putative III | - |
| GCA_008641105.1 | VTTIO1000003.1 128 | H. salina XCD-X85 | 291 | 134.6 | Low | Putative III | - |
| GCA_008641105.1 | VTTIO1000003.1 202 | H. salina XCD-X85 | 308 | 386.7 | High | Putative IIa | - |
| GCA_027474385.1 | CP114836.1 2415 | H. sp MFKK-1 | 289 | 142.3 | Low | Putative III | - |
| GCA_027474385.1 | CP114836.1 1279 | H. sp MFKK-1 | 304 | 395.2 | High | Putative IIa | - |
| GCA_021545785.1 | CP079801.1 395 | H. sp RR6 | 289 | 147.9 | Low | Putative III | - |
| GCA_021545785.1 | CP079801.1 1931 | H. sp RR6 | 305 | 387.3 | High | Putative IIa | - |
| GCA_021545785.1 | CP079801.1 1930 | H. sp RR6 | 304 | 387.6 | High | Putative IIa | - |
| GCA_029866945.1 | CP123445.1 1947 | H. sp SMJS2 | 287 | 140 | Low | Putative III | - |
| GCA_029866945.1 | CP123445.1 1693 | H. sp SMJS2 | 302 | 398.9 | High | Putative IIa | *PbauzCut*<sup>4</sup> |
| GCA_014219065.1 | JACLGO010000004.1 55 | H. xiamenensis PX1 | 285 | 148.2 | Low | Putative III | - |
| GCA_014219065.1 | JACLGO010000006.1 8 | H. xiamenensis PX1 | 302 | 403.8 | High | Putative IIa | - |
| GCA_011050525.1 | DRFO01000030.1 14 | H. xinjiangensis HyVt-324 | 287 | 139.2 | Low | Putative III | - |
| GCA_011050525.1 | DRFO01000030.1 15 | H. xinjiangensis HyVt-324 | 289 | 142.8 | Low | Putative III | - |
| GCA_011053075.1 | DRHP01000073.1 14 | H. xinjiangensis HyVt-372 | 286 | 132.5 | Low | Putative III | - |
| GCA_011053075.1 | DRHP01000073.1 15 | H. xinjiangensis HyVt-372 | 287 | 139.2 | Low | Putative III | - |
| GCA_011053075.1 | DRHP01000073.1 16 | H. xinjiangensis HyVt-372 | 289 | 142.8 | Low | Putative III | - |
| GCA_011050525.1 | DRFO01000030.1 13 | H. xinjiangensis HyVt-324 | 286 | 132.5 | Low | Putative III | - |
| GCA_900108005.1 | FNVE01000001.1 785 | P. aestusnigri CECT 8317 | 289 | 143.7 | Low | Putative III | - |
| GCA_900108005.1 | FNVE01000007.1 96 | P. aestusnigri CECT 8317 | 304 | 396.3 | High | Ila | Haes_PE-H<sup>3</sup> |
| GCA_900114735.1 | FOUA01000001.1 295 | P. bauzanensis CGMCC 1.9095 | 287 | 142.8 | Low | Putative III | - |
| GCA_900114735.1 | FOUA01000001.1 46 | P. bauzanensis CGMCC 1.9095 | 302 | 398.9 | High | Ila | PbauzCut<sup>4</sup> |
| GCA_900111225.1 | FOGN01000001.1 966 | P. bauzanensis DSM 22558 | 287 | 142.8 | Low | Putative III | - |
| GCA_900111225.1 | FOGN01000001.1 1216 | P. bauzanensis DSM 22558 | 302 | 398.9 | High | Ila | PbauzCut<sup>4</sup> |
| GCA_900115905.1 | FOYD01000002.1 336 | P. formosensis JCM 18415 | 287 | 144.4 | Low | Putative III | - |
| GCA_900115905.1 | FOYD01000004.1 228 | P. formosensis JCM 18415 | 304 | 395.1 | High | Putative Ila | - |
| GCA_900105005.1 | LT629748.1 1969 | P. litoralis 2SM5 | 287 | 145.1 | Low | Putative III | - |
| GCA_900105005.1 | LT629748.1 1727 | P. litoralis 2SM5 | 302 | 401.7 | High | Putative Ila | - |
| GCA_020025155.1 | CP073751.1 2021 | P. nanhaiensis SCS 2-3 | 287 | 126.5 | Low | Putative III | - |
| GCA_020025155.1 | CP073751.1 2020 | P. nanhaiensis SCS 2-3 | 286 | 137.7 | Low | Putative III | - |
| GCA_020025155.1 | CP073751.1 2019 | P. nanhaiensis SCS 2-3 | 287 | 154 | Low | Putative III | - |
| GCA_020025155.1 | CP073751.1 1339 | P. nanhaiensis SCS 2-3 | 300 | 387.2 | High | Putative Ila | - |
| GCA_020025155.1 | CP073751.1 1341 | P. nanhaiensis SCS 2-3 | 302 | 412.4 | High | Putative Ila | - |
| GCA_014641295.1 | BMHR01000028.1 10 | P. oceani CGMCC 1.15195 | 288 | 132.8 | Low | Putative III | - |
| GCA_014641295.1 | BMHR01000002.1 361 | P. oceani CGMCC 1.15195 | 303 | 395.2 | High | Putative Ila | - |
| GCA_900114765.1 | FOUD01000029.1 9 | P. pachastrellae JCM 12285 | 289 | 139.6 | Low | Putative III | - |
| GCA_900114765.1 | FOUD01000014.1 50 | P. pachastrellae JCM 12285 | 304 | 387 | High | Putative Ila | - |
| GCA_900105255.1 | LT629763.1 218 | P. sabulinigri JCM 14963 | 289 | 142.6 | Low | Putative III | - |
| GCA_900105255.1 | LT629763.1 219 | P. sabulinigri JCM 14963 | 289 | 146.2 | low | Putative III | - |
| GCA_900105255.1 | LT629763.1 2628 | P. sabulinigri JCM 14963 | 303 | 382 | High | Putative Ila | - |
| GCA_900105255.1 | LT629763.1 2629 | P. sabulinigri JCM 14963 | 305 | 383.4 | High | Putative Ila | - |
| GCA_900105655.1 | LT629787.1 958 | P. salegens CECT 8338 | 289 | 140.3 | Low | Putative III | - |
| GCA_900105655.1 | LT629787.1 957 | P. salegens CECT 8338 | 288 | 140.7 | Low | Putative III | - |
| GCA_900105655.1 | LT629787.1 2802 | P. salegens CECT 8338 | 315 | 383.5 | High | Putative IIa | - |
| GCA_900105655.1 | LT629787.1 2801 | P. salegens CECT 8338 | 315 | 392.2 | High | Putative IIa | - |
| GCA_014637955.1 | BMFF01000001.1 657 | P. salina CGMCC 1.12482 | 293 | 134.2 | Low | Putative III | - |
| GCA_014637955.1 | BMFF01000001.1 656 | P. salina CGMCC 1.12482 | 291 | 134.6 | Low | Putative III | - |
| GCA_014637955.1 | BMFF01000001.1 582 | P. salina CGMCC 1.12482 | 308 | 386.7 | High | Putative IIa | - |
| GCA_900104945.1 | LT629736.1 1619 | P. xinjiangensis NRRL B-51270 | 285 | 142.6 | Low | Putative III | - |
| GCA_900104945.1 | LT629736.1 1618 | P. xinjiangensis NRRL B-51270 | 287 | 151.6 | Low | Putative III | - |
| GCA_900104945.1 | LT629736.1 889 | P. xinjiangensis NRRL B-51270 | 303 | 405.3 | High | Putative IIa | - |
| GCA_900114825.1 | FOUI01000001.1 410 | P. yangmingensis DSM 24213 | 286 | 134.6 | Low | Putative III | - |
| GCA_900114825.1 | FOUI01000004.1 1 | P. yangmingensis DSM 24213 | 303 | 380.7 | High | Putative IIa | - |
| GCA_900104945.1 | LT629736.1 1620 | P.xinjiangensis NRRL B-51270 | 288 | 125.2 | Low | Putative III | - |

**Table S5:**
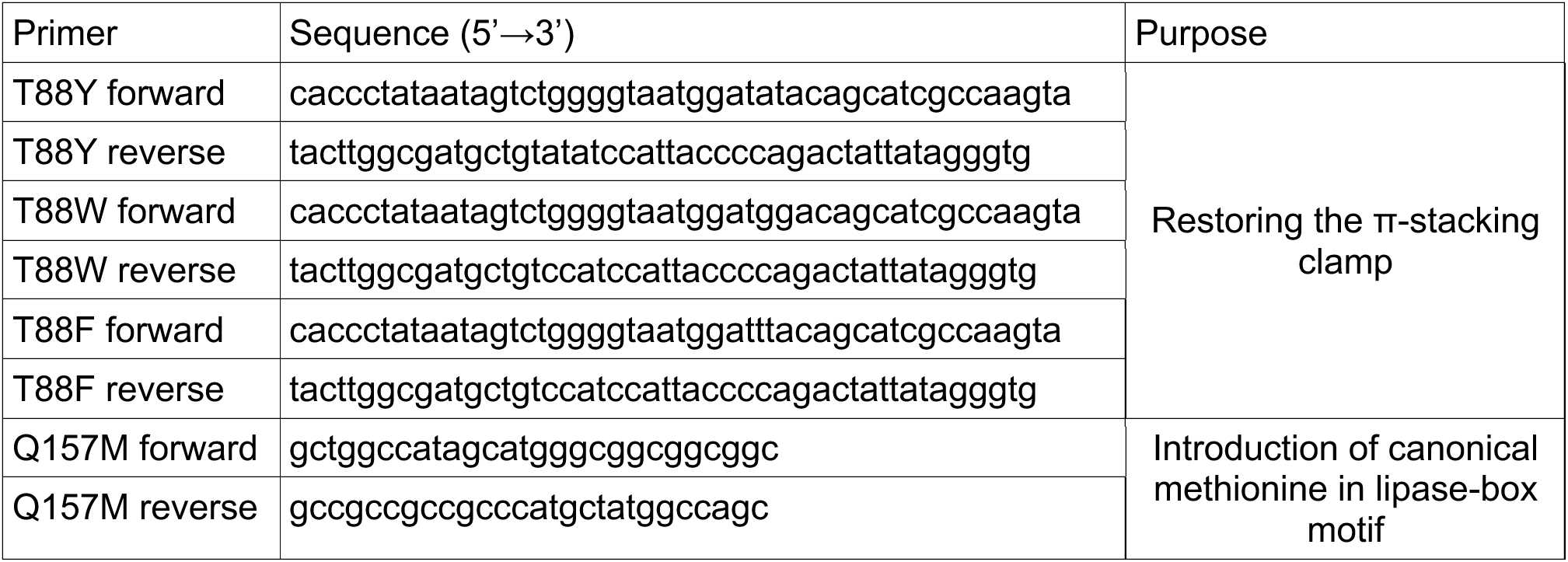
Primer pairs for the introduction of single mutations in *Halo*PETase-1 by site-directed mutagenesis.

## References

1. 1. Environment, U. N. Turning off the Tap: How the world can end plastic pollution and create a circular economy | UNEP - UN Environment Programme. https://www.unep.org/resources/turning-off-tap-end-plastic-pollution-create-circular-economy (2023).

2. 2. End plastic pollution: Towards an international legally binding instrument. https://wedocs.unep.org/bitstream/handle/20.500.11822/38525/k2200647_-_unep-ea-5-l-23-rev-1_-_advance.pdf?sequence=1&isAllowed=y (2022).

3. Geyer, R., Jambeck, J. & Law, K. L. Production, use, and fate of all plastics ever made. Science Advances 3, (2017).

4. 4. Ritchie, Hannah, Samborska, Veronika & Roser, Max. Plastic Pollution. https://ourworldindata.org/plastic-pollution (2023).

5. Cui, Y. et al. Computational redesign of a hydrolase for nearly complete PET depolymerization at industrially relevant high-solids loading. Nat Commun 15, 1417 (2024).

6. Tournier, V. et al. An engineered PET depolymerase to break down and recycle plastic bottles. Nature 580, 216–219 (2020).

7. Pfaff, L. et al. Multiple Substrate Binding Mode-Guided Engineering of a Thermophilic PET Hydrolase. ACS Catal. 12, 9790–9800 (2022).

8. Lu, H. et al. Machine learning-aided engineering of hydrolases for PET depolymerization. Nature 604, 662–667 (2022).

9. Sadler, J. C. & Wallace, S. Microbial synthesis of vanillin from waste poly(ethylene terephthalate). Green Chem. 23, 4665–4672 (2021).

10. Carniel, A., Waldow, V. de A. & Castro, A. M. de. A comprehensive and critical review on key elements to implement enzymatic PET depolymerization for recycling purposes. Biotechnology Advances 52, 107811 (2021).

11. Sonnendecker, C. et al. Low Carbon Footprint Recycling of Post-Consumer PET Plastic with a Metagenomic Polyester Hydrolase. ChemSusChem 15, (2022).

12. Richter, P. K. et al. Structure and function of the metagenomic plastic-degrading polyester hydrolase PHL7 bound to its product. Nat Commun 14, 1905 (2023).

13. Arnal, G. et al. Assessment of Four Engineered PET Degrading Enzymes Considering Large- Scale Industrial Applications. ACS Catal. 13, 13156–13166 (2023).

14. Fritzsche, S., Tischer, F., Peukert, W. & Castiglione, K. You get what you screen for: a benchmark analysis of leaf branch compost cutinase variants for polyethylene terephthalate (PET) degradation. *React*. Chem. Eng. 8, 2156–2169 (2023).

15. Thomsen, T. B., Hunt, C. J. & Meyer, A. S. Influence of substrate crystallinity and glass transition temperature on enzymatic degradation of polyethylene terephthalate (PET). New Biotechnology 69, 28–35 (2022).

16. Thomsen, T. B., Radmer, T. S. & Meyer, A. S. Enzymatic degradation of poly(ethylene terephthalate) (PET): Identifying the cause of the hypersensitive enzyme kinetic response to increased PET crystallinity. Enzyme and Microbial Technology 173, 110353 (2024).

17. Uekert, T. et al. Life cycle assessment of enzymatic poly(ethylene terephthalate) recycling. Green Chem. 24, 6531–6543 (2022).

18. Singh, A. et al. Techno-economic, life-cycle, and socioeconomic impact analysis of enzymatic recycling of poly(ethylene terephthalate). Joule 5, 2479–2503 (2021).

19. Uekert, T. et al. Technical, Economic, and Environmental Comparison of Closed-Loop Recycling Technologies for Common Plastics. ACS Sustainable Chem. Eng. 11, 965–978 (2023).

20. Sunagawa, S. et al. Structure and function of the global ocean microbiome. Science 348, 1261359 (2015).

21. Sana, B., Ghosh, D., Saha, M. & Mukherjee, J. Purification and characterization of an extremely dimethylsulfoxide tolerant esterase from a salt-tolerant *Bacillus* species isolated from the marine environment of the *Sundarbans*. Process Biochemistry 42, 1571–1578 (2007).

22. Wu, G. et al. A cold-adapted, solvent and salt tolerant esterase from marine bacterium *Psychrobacter pacificensis*. International Journal of Biological Macromolecules 81, 180–187 (2015).

23. Yang, Q. et al. Characterization of a Novel, Cold-Adapted, and Thermostable Laccase-Like Enzyme With High Tolerance for Organic Solvents and Salt and Potent Dye Decolorization Ability, Derived From a Marine Metagenomic Library. Front. Microbiol. 9, (2018).

24. Ghattavi, S. & Homaei, A. Marine enzymes: Classification and application in various industries. International Journal of Biological Macromolecules 230, 123136 (2023).

25. Chen, J. et al. Global marine microbial diversity and its potential in bioprospecting. Nature 633, 371–379 (2024).

26. Karan, R., Capes, M. D. & DasSarma, S. Function and biotechnology of extremophilic enzymes in low water activity. Aquatic Biosystems 8, 4 (2012).

27. Sun, J. et al. Solubilization and Upgrading of High Polyethylene Terephthalate Loadings in a Low-Costing Bifunctional Ionic Liquid. ChemSusChem 11, 781–792 (2018).

28. Xu, A., Zhou, J., Blank, L. M. & Jiang, M. Future focuses of enzymatic plastic degradation. Trends in Microbiology 31, 668–671 (2023).

29. Buchholz, P. C. F. et al. Plastics degradation by hydrolytic enzymes: The plastics-active enzymes database—PAZy. Proteins: Structure, Function, and Bioinformatics 90, 1443–1456 (2022).

30. Danso, D. et al. New Insights into the Function and Global Distribution of Polyethylene Terephthalate (PET)-Degrading Bacteria and Enzymes in Marine and Terrestrial Metagenomes. Applied and Environmental Microbiology 84, e02773–17 (2018).

31. Erickson, E. et al. Sourcing thermotolerant poly(ethylene terephthalate) hydrolase scaffolds from natural diversity. Nat Commun 13, 7850 (2022).

32. Perez-Garcia, P. et al. An archaeal lid-containing feruloyl esterase degrades polyethylene terephthalate. Commun Chem 6, 1–13 (2023).

33. Joo, S. et al. Structural insight into molecular mechanism of poly(ethylene terephthalate) degradation. Nature Communications 9, 382–382 (2018).

34. Bollinger, A. et al. A Novel Polyester Hydrolase From the Marine Bacterium Pseudomonas aestusnigri – Structural and Functional Insights. Front. Microbiol. 11, 114 (2020).

35. Rudra, B. & Gupta, R. S. Phylogenomic and comparative genomic analyses of species of the family Pseudomonadaceae: Proposals for the genera Halopseudomonas gen. nov. and Atopomonas gen. nov., merger of the genus Oblitimonas with the genus Thiopseudomonas, and transfer of some misclassified species of the genus Pseudomonas into other genera. International Journal of Systematic and Evolutionary Microbiology 71, 005011 (2021).

36. Bollinger, A., Thies, S., Katzke, N. & Jaeger, K.-E. The biotechnological potential of marine bacteria in the novel lineage of Pseudomonas pertucinogena. Microbial Biotechnology 13, 19–31 (2020).

37. de Witt, J. et al. Biodegradation of poly(ester-urethane) coatings by Halopseudomonas formosensis. Microbial Biotechnology 17, e14362 (2024).

38. Avilan, L. et al. Concentration-Dependent Inhibition of Mesophilic PETases on Poly(ethylene terephthalate) Can Be Eliminated by Enzyme Engineering. ChemSusChem 16, e202202277 (2023).

39. Ronkvist, Å. M., Xie, W., Lu, W. & Gross, R. A. Cutinase-Catalyzed Hydrolysis of Poly(ethylene terephthalate). Macromolecules 42, 5128–5138 (2009).

40. Weigert, S., Gagsteiger, A., Menzel, T. & Höcker, B. A versatile assay platform for enzymatic poly(ethylene-terephthalate) degradation. *Protein Engineering*, Design and Selection 34, gzab022 (2021).

41. Schmidt, J. et al. Effect of Tris, MOPS, and phosphate buffers on the hydrolysis of polyethylene terephthalate films by polyester hydrolases. FEBS Open Bio 6, 919–927 (2016).

42. Thomsen, T. B. et al. Rate Response of Poly(Ethylene Terephthalate)-Hydrolases to Substrate Crystallinity: Basis for Understanding the Lag Phase. ChemSusChem 16, e202300291 (2023).

43. Chen, C.-C. et al. General features to enhance enzymatic activity of poly(ethylene terephthalate) hydrolysis. Nat Catal 4, 425–430 (2021).

44. Burgin, T. et al. The reaction mechanism of the Ideonella sakaiensis PETase enzyme. Commun Chem 7, 65 (2024).

45. Mirdita, M. et al. ColabFold: making protein folding accessible to all. Nat Methods 19, 679– 682 (2022).

46. Yoshida, S. et al. A bacterium that degrades and assimilates poly(ethylene terephthalate). Science 351, 1196–1199 (2016).

47. Genome- and Community-Level Interaction Insights into Carbon Utilization and Element Cycling Functions of Hydrothermarchaeota in Hydrothermal Sediment | mSystems. https://journals.asm.org/doi/10.1128/msystems.00795-19.

48. Eddy, S. R. Profile hidden Markov models. Bioinformatics 14, 755–763 (1998).

49. Teufel, F. et al. SignalP 6.0 predicts all five types of signal peptides using protein language models. Nat Biotechnol 40, 1023–1025 (2022).

50. Geertsma, E. R. & Dutzler, R. A Versatile and Efficient High-Throughput Cloning Tool for Structural Biology. Biochemistry 50, 3272–3278 (2011).

51. Bjerga, G. E. K., Arsın, H., Larsen, Ø., Puntervoll, P. & Kleivdal, H. T. A rapid solubility- optimized screening procedure for recombinant subtilisins in E. coli. Journal of Biotechnology 222, 38–46 (2016).

52. Pérez-García, P., Danso, D., Zhang, H., Chow, J. & Streit, W. R. Exploring the global metagenome for plastic-degrading enzymes. in Methods in Enzymology vol. 648 137–157 (Elsevier, 2021).

53. Hyatt, D. et al. Prodigal: prokaryotic gene recognition and translation initiation site identification. BMC Bioinformatics 11, 119 (2010).

54. Cianci, M. et al. P13, the EMBL macromolecular crystallography beamline at the low- emittance PETRA III ring for high- and low-energy phasing with variable beam focusing. J Synchrotron Rad 24, 323–332 (2017).

55. Krug, M., Weiss, M. S., Heinemann, U. & Mueller, U. XDSAPP: a graphical user interface for the convenient processing of diffraction data using XDS. J Appl Cryst 45, 568–572 (2012).

56. Adams, P. D. et al. PHENIX: a comprehensive Python-based system for macromolecular structure solution. Acta Cryst D 66, 213–221 (2010).

57. Emsley, P., Lohkamp, B., Scott, W. G. & Cowtan, K. Features and development of Coot. Acta Crystallogr D Biol Crystallogr 66, 486–501 (2010).

58. Berman, H. M. et al. The Protein Data Bank. Nucleic Acids Research 28, 235–242 (2000).

59. The PyMOL Molecular Graphics System. Schrödinger, LLC (2024).

60. Do, C. B., Mahabhashyam, M. S. P., Brudno, M. & Batzoglou, S. ProbCons: Probabilistic consistency-based multiple sequence alignment. Genome Res. 15, 330–340 (2005).

61. The UniProt Consortium. UniProt: the Universal Protein Knowledgebase in 2023. Nucleic Acids Research 51, D523–D531 (2023).

62. Larkin, M. A. et al. Clustal W and Clustal X version 2.0. Bioinformatics 23, 2947–2948 (2007).

63. Zhu, L. Handbook of Thermal Analysis and Calorimetry Applications to Polymers and Plastics. (Elsevier Science B.V., 2002).

## Supplementary references

1. Benson, D. A. et al. GenBank. Nucleic Acids Research 41, D36–D42 (2013).

2. Altschul, S. F., Gish, W., Miller, W., Myers, E. W. & Lipman, D. J. Basic local alignment search tool. Journal of Molecular Biology 215, 403–410 (1990).

3. Bollinger, A. et al. A Novel Polyester Hydrolase From the Marine Bacterium Pseudomonas aestusnigri – Structural and Functional Insights. Front. Microbiol. 11, 114 (2020).

4. Avilan, L. et al. Concentration-Dependent Inhibition of Mesophilic PETases on Poly(ethylene terephthalate) Can Be Eliminated by Enzyme Engineering. ChemSusChem 16, e202202277 (2023).

5. Larkin, M. A. et al. Clustal W and Clustal X version 2.0. Bioinformatics 23, 2947–2948 (2007).

6. Gonnet, G. H., Cohen, M. A. & Benner, S. A. Exhaustive Matching of the Entire Protein Sequence Database. Science 256, 1443–1445 (1992).

7. Joo, S. et al. Structural insight into molecular mechanism of poly(ethylene terephthalate) degradation. Nature Communications 9, 382–382 (2018).

